# Iron regulates contrasting toxicity of uropathogenic *Eschericia coli* in macrophages and epithelial cells

**DOI:** 10.1101/2022.05.29.493834

**Authors:** Deepti Dabral, Hiren Ghosh, Masato Niwa, Tasuku Hirayama, Rinse de Boer, Marjon de Vos, Geert van den Bogaart

## Abstract

By far most urinary tract infections are caused by genetically diverse uropathogenic *Escherichia coli* (UPEC). Knowledge of the virulence mechanisms of UPEC is critical for drug development, but most studies focus on only a single strain of UPEC. In this study, we compared the virulence mechanisms of four antibiotic-resistant and highly pathogenic UPEC isolates in human blood monocyte-derived macrophages and a bladder epithelial cell (BEC) line: ST999, ST131, ST1981 and ST95. We found that while non-pathogenic *E. coli* strains are efficiently killed by macrophages in bactericidal single membrane vacuoles, the UPEC strains survive within double-membrane vacuoles. On side-by-side comparison, we found that whereas ST999 only carries Fe^3+^ importers, ST95 carries both Fe^2+^ and Fe^3+^ importers and the toxins haemolysin and colibactin. Moreover, we found that ST999 grows in the Fe^3+^ rich vacuoles of BECs and macrophages with concomitant increased expression of haem receptor *chuA* and the hydrogen peroxide sensor *oxyR*. In contrast, ST95 produces toxins in iron-depleted conditions similar to that of the urinary tract. Whereas ST95 also persist in the iron rich vacuoles of BECs, it produces colibactin in response to low Fe^3+^ contributing to macrophage death. Thus, iron regulates the contrasting toxicities of UPEC strains in macrophages and bladder epithelial cells due to low and high labile iron concentrations, respectively.

**Key findings:**

- Antibiotics resistant uropathogenic *E. coli* strains ST999, ST131, ST1981, and ST95 survive within spacious double membrane vacuoles. Non-pathogenic *E. coli* strains XL1 blue and MG1655 are cleared in single membrane vacuoles in macrophages.
- ST999 lacks Fe^2+^ importer and toxins, and grows in iron rich vacuoles of macrophages and bladder epithelial cells.
- ST95 carries both Fe^2+^ and Fe^3+^ importers and grows in iron low conditions.
- ST95 expresses toxins and induces cell death of infected macrophages, but not of bladder epithelial cells.
- Bladder epithelial cells have a higher pool of labile iron than macrophages. Differential expression of virulence factors by ST999 and ST95 in bladder epithelial cells and macrophages is dependent on iron concentration.

## Introduction

In all humans, *Escherichia coli* inhabits the gastrointestinal tract, where it is part of the healthy commensal microbiota. However, certain pathogenic strains, called uropathogenic *E. coli* (UPEC), are derived from this commensal *E. coli* but account for >80% urinary tract infections (UTIs) [1-3]. These are the most common infections, as >50% of women suffer from it at least once during their lifetime [4]. UTIs are often difficult to treat due to the formation of intracellular *E. coli* colonies within bladder epithelial cells (BECs) which confer antibiotic resistance [2, 5, 6]. These hidden colonies are a source of recurrent infections in 20-30% of patients [7]. UPEC evolved to survive in two very different environments: the gastrointestinal and the urinary tract. Because of the low nutrient contents, especially iron with a mean concentration of 101 ± 45 μg/L in urine of healthy adults [8], and high concentrations of the protein denaturing agent urea (∼23 mM) [3], the urinary tract is a hostile environment for *E. coli*. To counter these challenges, UPEC evolved mechanisms to survive in this environment [9, 10]. Importantly, UPEC can invade BECs and proliferate within the nutrient-rich intracellular environment, and this niche protects it from the surveilling macrophages and other immune cells [11, 12]. Tissue-resident macrophages are among the first immune cells to respond to infected bladder tissue [4, 13] and comprise ∼40% of all immune cells in the murine bladder [14]. Infected BECs secrete CCX2 chemokine for the recruitment of CD14+ monocytes from blood, which also differentiate into macrophages and supplement the tissue-resident macrophages [13, 15]. At the infection site, macrophages clear UPEC by phagocytosis and cytotoxicity mechanisms [16, 17].

Clinical isolates of UPEC are known to be genetically diverse [18] but how this affects the virulence mechanisms is incompletely understood. UPEC isolates can be genotyped based on sequence variations in certain genes, and categorized in phylogeny. In multi-locus sequence typing (MLST), nucleotide sequence variations are measured within seven housekeeping genes (*adk, fumC, gyrB, icd, mdh, purA* and *recA*) to assign sequence types (ST) to *E. coli* strains [19, 20]. Major pandemic clonal lineages of UPEC are ST131, ST393, ST69, ST95, and ST73 [20]. Other typing methods are based on polymorphisms in the O antigen, which is the outer polysaccharides of lipopolysaccharide (LPS), the H antigen of flagella, and the *fimH* gene involved in fimbriae [21, 22]. As a consequence of genetic differences, UPEC strains carry functionally redundant iron acquisition genes [23] and toxins (*senB, clb, hly, cnf1*) [24-26]. Haemolysin (*hly*) mediates formation of pores in the membrane of host cells, and both colibactin (*clb*) and Shigella enterotoxin (*senB*) cause DNA damage to the host cell leading to cell death [26-28]. As a consequence of genetic diversity, UPEC strains can be expected to have different virulence mechanisms.

However, most studies addressing the virulence mechanisms of UPEC use a single strain, mostly UTI89 (typed as ST95) and CFT073 (typed as ST73) [29-31]. In this study, we performed whole genome sequencing for genotyping, identification of antibiotic resistant genes of four UPEC isolates. These are typed as ST999 (*fimH20; O null: H4*), ST131 (*fimH30; O25:H4*), ST1981 (*fimH49; O1:H6*), and ST95 (fimH18; *O18:H7*). We found that all these UPEC strains survive in LAMP1 positive double-membrane vacuoles in macrophages, whereas non-pathogenic *E. coli* strains are efficiently killed in single-membrane vacuoles. With detailed side-by-side comparison of ST999 and ST95, we show that ST95 is better adapted to grow in low extracellular iron concentrations of the urinary tract, as it carries high affinity iron acquisition genes. Moreover, genes encoding toxins were expressed at higher levels in iron depleted conditions by ST95. These toxins likely contribute to the selective killing by ST95 of macrophages, but not of BECs which have a higher content of labile iron. In contrast, we found that ST999 is better adapted to grow at higher iron concentrations and survived intracellularly in both BECs and macrophages. Thus, the virulence mechanisms of UPEC differ depending on the strain and host cell type.

## Material and Methods

### Bacterial DNA isolation

UPEC strains were cultured overnight in LB broth media at 37°C. Bacteria culture (0.5 ml) was centrifuged at 2000 ×g for 5 mins to pellet cells. After PBS wash, cells were lysed in 0.5 ml solution containing 20% sucrose, 10 mM Tris-HCl, pH 8.1, 10 mM EDTA, 50 mM NaCl, 5 mg/ml lysozyme, and 100 μg/ml RNaseA at 55°C for 10 mins. After 10 mins, 25 μl of 20 mg/ml proteinase K and 10% SDS were added and bacteria were incubated for 1 h at 60°C. Total DNA was extracted using phenol/chloroform/isoamyl alcohol (25:24:1; *v*/*v*/*v*). The upper aqueous layer containing total DNA was collected and precipitated by centrifugation at 12,0000 ×g in the presence of 3 M sodium acetate and isopropyl alcohol. DNA pellet was washed with 70% ethanol and dissolved in Tris-EDTA buffer.

### Whole genome library construction and sequencing

Whole genome sequencing was carried out on the MGISEQ-2000 platform at BGI (Shenzhen, Denmark). The insert size of the library was 350 bp with a pair-end sequencing length of 150 bp. Briefly, 1 µg genomic DNA was randomly fragmented by a g-TUBE device (Covaris, Inc). The fragmented DNA was selected by magnetic beads to an average size of 200 - 400 bp. The selected fragments were end repaired, 3’ adenylated, adapters-ligated, PCR amplified and the products were purified by the magnetic beads. The stranded PCR products were heat denatured and circularized by the splint oligo sequence. The single stranded circular DNA was formatted as the final library.

### Read processing, genome assembly and annotation

Sequence reads were assembled using SPAdes v3.11.1 [32] with default parameter. Assemblies were filtered that included contigs size ≥500 bp and annotated with Prokka v1.14.5 [33]. Multi-locus sequence typing (MLST) was carried out with mlst v2.10 [34]. Antibiotic resistance gene, and virulence genes were identified using Abricate v0.9.8 against the database Resfinder and E. coli_vf, respectively with the default settings (https://github.com/tseemann/abricate). Identification of phylogroup and serotype was carried out using Clermon Typing v1.4.0 [35], and SerotypeFinder v2.0 [36], respectively. Prophages were identified using Phaster webserver [37]. A pan genome matrix and core genome alignment were generated by Roary v.3.13.0 [38] with default parameters. For core genome phylogenetic reconstructions Fastree [39] was used. Genomic synteny comparison was performed using BLAST Ring Image Generator (BRIG) [40].

### Mammalian cell culture

Approval to conduct experiments with human blood samples was obtained from the blood bank and all experiments were conducted according to national and institutional guidelines. Informed consent was obtained from all blood donors by the Dutch blood bank Sanquin (Amsterdam, Netherlands). Samples were anonymized and none of the investigators could ascertain the identity of the donors. CD14+ monocytes were derived from buffy coats of healthy donors using standardized Ficoll method [41]. Briefly, PBS containing 2 mM EDTA was added to the buffy coat, followed by centrifugation to collect the peripheral blood mononuclear cells. After washing, CD14+ microbeads (Miltenyi Biotec, cat.no: 130-097-052) and magnetic antibody cell sorting (MACS) was used according to manufacturer’s protocol. Monocytes were seeded in low attachment plates (Corning Costar, cat.no: 3471) to differentiate at 37°C, 5% CO^2^ for 8 days with 100 ng/ml macrophage colony-stimulating factor (M-CSF; R&D systems, cat.no 216-MC).

Human bladder carcinoma 5637 cells (ATCC, HTB-9) were grown in RPMI-1640 media (GIBCO, cat.no. 21875-034) containing 10% fetal bovine serum (FBS) (Fisher Scientific, cat. no. 11521831).

### Bacteria culture

ST999, ST131, ST1981, ST95, *E. coli* XL 1 blue and K-12 MG1655 were cultured overnight in Luria Bertani (LB) broth media (Sigma Aldrich, cat. no: L3022). Overnight cultures were diluted to OD 600 ∼0.5. Bacterial suspension was further diluted to yield required bacteria for infection. All infections were carried at the ratio of 10 macrophages per bacterium and 300 BECs per bacterium unless stated otherwise.

### Growth curves

Overnight cultures in LB broth media were washed with PBS and adjusted to ∼0.5 OD 600 nm in M9 broth media (Fluka, cat. no. 63011) supplemented with 1 M MgSO^4^ and 20% glucose. Bacteria suspension (10 μl) was inoculated to 100 μl of M9 media, M9 media was supplemented with 0.1 mM FeCl^3^ (Sigma Aldrich, cat. no. 44944) or with 1 μM deferoxamine (Sigma Aldrich, cat. no. D9533) in a 96-well plate (Costar, cat no. 3370). OD 600 nm measurements were carried out using a BioTek Synergy HTX plate reader.

### Colony forming unit (CFU) assay

Infections of ∼15,000 macrophages were carried out at the ratio of 50 macrophages per bacterium. BECs were grown to 100% confluent monolayers yielding 470,000 ± 6,700 cells per well. Infection of BECs were carried out at the ratio of 1,500 BECs per bacterium. After 1 h, RPMI-1640 media containing 10% FBS was supplemented with antibiotics (6 mg/l Nitrofurantoin, Sigma Aldrich, cat. no. N7878; 16 mg/l Imipenem monohydrate, Sigma Aldrich, cat. no. I0160; 200 mg/l; Gentamicin, Fisher Scientific, cat. no 12664735; 25 mg/l Trimethoprim, Sigma Aldrich, cat. no. 92131; 125 mg/l sulfamethoxazole, Sigma Aldrich, cat. no. S7507) to kill extracellular bacteria. Another antibiotic treatment for 2 h was also carried out before lysis. At 4 and 6 h post-infection (hpi), macrophages and BECs were washed and lysed using 0.1% Triton-X100. Lysates were cultured on LB agar plates at 37°C overnight.

### Flow cytometry

150,000 macrophages were seeded in low attachment 6 well plates (Corning Costar, cat. no. CLS7007-24EA). Infections were carried out at the ratio of 10 macrophages per bacterium. At 1 h, macrophages were stained with Apoptosis Detection Kit (BD Biosciences; cat. no: 556570), and analyzed by Flow cytometry (CytoFlexS, Beckman Coulter) using appropriate filters.

### Lactate dehydrogenase (LDH) assay

BECs were cultured to 100% confluence yielding 470,000 ± 6,700 BECs per well. Infections were carried out at the ratio of 10 BECs per bacterium. >1% DMSO was added to the control wells. At 1 hpi, media was collected and LDH activity measurement was carried out in 96-well plate (Costar, cat no. 3370) according to LDH assay kit (Abcam, cat. no ab102526). Absorbance measurement at 450 and 570 nm was carried out using microplate reader (BioTek, Synergy HTX).

### Enzyme linked immunosorbent assay (ELISA)

15, 000 macrophages were infected at the ratio of 10 macrophages per bacterium and 470,000 ± 6,700 BECs were infected at the ratio of 300 BECs per bacterium. Media was collected at 4 and 6 hpi and filtered through 0.2 μm filter to remove dead bacteria and cellular debris. IL-6 (Invitrogen; cat. no 88-7064) and TNF-α (Invitrogen; cat. no 88-7346) ELISAs were carried out according to the manufacturer’s instructions. Absorbance measurements at 450 and 570 nm were carried out using a microplate reader (BioTek, Synergy HTX).

### Hemolysin assessment

15, 000 macrophages and 470,000 ± 6,700 BECs were infected at the ratio of 10 macrophages per bacterium and 300 BECs per bacterium, respectively. ST999, ST131, ST1981 and ST95 infected macrophages and BECs were lysed at 1 hpi using 0.1% Triton X-100. Lysate was cultured overnight at 37°C on agar medium containing sheep blood (Fisher Scientific, cat no. PB5012A).

### Labile iron concentration

500 macrophages and BECs were seeded in 96-well plate (Costar, cat no. 3370) in triplicates. Cells were incubated with 1 μM calcein-AM (Biotium, cat no. 80011-3) for 10 mins. The chelatable pool was determined by incubating cells with 30 μM pyridoxal isonicotinoyl hydrazone (PIH) (Abcam, cat no. ab145871) for 10 mins, prior to incubation with calcein-AM. 0.1% DMSO was added to the control wells to subtract background measurements. Fluorescence measurements with Ex/Em 485/528 nm were carried out using a microplate reader (BioTek, Synergy HTX).

### Immunofluorescence labeling and microscopy

15,000 macrophages and 470,000 ± 6,700 BECs were cultured on sterile coverslips of 12 mm diameter (Electron Microscopy Sciences, cat.no.72230-01). Macrophages were infected at the ratio of 10 macrophages per bacterium and BECs were infected at the ratio of 300 BECs per bacterium. At 4 hpi, macrophages and BECs were fixed with 4% paraformaldehyde (Aurion, cat. no. 15710) and immuno stained for LAMP1 (Sigma Aldrich, cat no. L1418), LPS (Invitrogen, cat. no. PA1-73178) and LC3B (Cell Signaling, cat no. 83506S). Phalloidin AF488 (Invitrogen, cat. no. A12379) was used to stain F-actin and DAPI (Sigma Aldrich, cat. no. 32670) stained nucleic acid. Image acquisition was done with a confocal laser scanning microscope (Zeiss, LSM 800) equipped with 64× oil immersion objective lens.

### Electron microscopy

15,000 macrophages were seeded on sterile coverslips of 12 mm diameter (Electron Microscopy Sciences, cat.no.72230-01). Infections were carried out at the ratio of 10 macrophages per bacterium. At 1 hpi, macrophages were fixed with 2% glutaraldehyde for 1 h in 0.1 M sodium cacodylate buffer pH 7.2. Macrophages are washed with sodium cacodylate and post fixed in 1% OsO^4^ in sodium cacodylate for 1 h. Samples were *en bloc* contrasted with 0.5% uranyl acetate overnight. Samples were dehydration in increasing concentration of ethanol and embedded in Epon resin. 80 nm sections were collected on formvar coated carbon evaporated copper grids and images were acquired using a CM12 transmission electron microscope (Philips) at 100 kV. Distances between the outer bacterial membrane and inner phagosomal membrane were measured at ≥5 positions per phagosome using ImageJ.

For electron tomography, 180 nm sections were collected on formvar coated carbon evaporated copper grids. Sections were decorated with 10 nm gold. Dual tilt tomograms were manually recorded including a tilt range of -45° to 45° with 2.5° interval with a CM12 transmission electron microscope running at 100 kV. Tomograms were reconstructed and 3D rendered using the Imod software.

### Iron probe labeling and live cell microscopy

900,000 macrophages were cultured in RPMI-1640 media (GIBCO, cat.no. 21875-034) containing 10% fetal bovine serum (FBS) (Fisher Scientific, cat. no. 11521831) in glass bottom culture dish (Greiner bio one, cat no. 627870). Macrophages were washed with PBS and maintained in RPMI-1640 without phenol red and glutamine (Gibco, cat no. 32404-014) and labelled with 1 μM Mem-RhoNox for 10 mins [42]. Excess probe was washed. Macrophages were maintained in RPMI-1640 without phenol red and glutamine for live cell microscopy.

Overnight culture of bacteria was washed with PBS and adjusted to ∼0.5 OD 600 nm for labelling with Alexa fluor 633 C5 maleimide for 30 mins. Excess probe was washed away. Infection of macrophages were carried out at the ratio of 10 macrophages per bacteria. Live cell microscopy was carried out using epi-fluorescence microscopy equipped with a LED excitation lamp, prime bSI sCMOS camera, Olympus 60× uAPO NA 1.49 oil-immersion objective lens, and appropriate filters.

### Western blotting

300,000 macrophages were infected at the ratio of 10 macrophages per bacterium. BECs were cultured to 100% confluence yielding 470,000 ± 6,700 BECs per well. Infections were carried out at the ratio of 300 BECs per bacterium. At 4 hpi, macrophages and BECs were lysed by boiling at 95° C for 5 mins in buffer containing 65.8 mM Tris HCl pH 6.8, 2.1 % SDS, 26.3 % (w/v) glycerol. Total proteins were quantified using EZQ protein quantification kit (Invitrogen, cat. no: R33200). Equal amounts of proteins were resolved using 4-20% SDS-PAGE (Bio-Rad, cat. no: 456-1094), and blotted onto 0.2 μm PVDF membranes (Bio-Rad, cat no. 1620177). Blots were incubated overnight at 4°C with following primary antibodies: LC3B (Novus Biologicals, cat. no. NB600-1384), uroplakin Ia (Abcam, cat. no. ab185970), and GAPDH (Cell signaling, cat. no. 2118). Secondary antibody was AF680 conjugated Goat anti-rabbit (Invitrogen, cat. no. A21076). Imaging was done using an Odyssey FC imager.

### Reverse transcriptase (RT) PCR

Overnight bacterial cultures in LB broth media were washed with PBS and adjusted to ∼0.5 OD 600 nm in M9 broth media (Fluka, cat. no. 63011) supplemented with 1 M MgSO^4^ and 20% glucose. Bacteria suspension (100 μl) was inoculated to 1 ml of M9 media, M9 media was supplemented with 0.1 mM FeCl^3^ (Sigma Aldrich, cat. no. 44944) or with 1 μM deferoxamine (Sigma Aldrich, cat. no. D9533). Cultures were allowed to grow until mid-log phase (8 h). Bacterial cells were washed and lysed by heating for 5 mins at 95°C with Max Bacterial Enhancement Reagent (Invitrogen, cat. no. 46-6036), and stored overnight at - 20°C.

For bacterial RNA isolation from infected BECs and macrophages, ≥ 300,000 macrophages and ∼470,000 ± 6,700 BECs were cultured. Macrophages and BECs were infected at the ratio of 10 macrophages per bacterium and 300 BECs per bacterium, respectively. At 1 hpi, macrophages and BECs were washed with PBS, and lysed by heating for 5 mins at 95°C with Max Bacterial Enhancement Reagent (Invitrogen, cat. no. 46-6036), and stored overnight at -20°C.

Total RNA was isolated by incubating bacterial lysates with Trizol reagent (Invitrogen, cat. no. 15596026) for 10 min, followed by extraction using phenol/chloroform. RNA concentrations were measured and cDNA was synthesized using random primers (Roche cat. no. 11034731001) and M-MLV reverse transcriptase (Invitrogen cat. no 28025-021). Target of interest was amplified using PowerUp SYBR Green Master mix (Applied Biosystems, cat no. A25741) and gene specific primers: *oxyR* (5’-GCTATCTATGAAGATCACCCGT-3’ and 5’-GTGACCATCTTCCAGCATCAG-3’), *chuA* (5’-CGATGGGCGAAATGTATAA-3’ and 5’-GTTAGTTTCCGGACGTAAG-3’), *iroN* (5’-AAGTCAAAGCAGGGGTTGCCCG-3’ and 5’-GACGCCGACATTAAGACGCAG-3’), *fliC* (5’-CCACGACAGGTCTTTATGATCTGA-3’ and 5’-CAACTGTGACTTTATCGCCATTCC-3’), *hma* (5’-ATCGTTCGGCAAGCAACCTTTG-3’ and 5’-ATGCGGATTTGTTTACGGCCTG-3’), *senB* (5’-CCGTTGAAAGATCCGAGACC-3’ and 5’-GTTTGGGTAGACCGGCATGT-3’), *clbN* (5’-GTTTTGCTCGCCAGATAGTCATTC-3’ and 5’-CAGTTCGGGTATGTGTGGAAGG-3’), *fimH (*5’-TGCAGAACGGATAAGCCGTGG-3’ and 5’-GCAGTCACCTGCCCTCCGGTA-3’), *16s* (5’-GGAGACTGCCAGTGATAAA-3’ and 5’-GAGGTCGCTTCTCTTTGT-3’). Amplification was carried out using a Bio-Rad CFX96 C1000 Thermal Cycler Real-Time System. Samples (*n* ≥ 3 biological replicates) were included in triplicate for each primer. Fold expression of the transcript, relative to 16s transcript is expressed as 2^ΔΔCt [43].

### Statistical analysis

Statistical analyses were performed using GraphPad Prism 5. Two tailed unpaired t-tests and one-way ANOVA was used for multiple comparisons. P-values <0.05 were considered significant. Data is reported as mean ± SEM.

## Results

### Antibiotic resistant UPEC strains are genetically diverse

Based on MLST, the UPEC strains used in this study were typed as ST999, ST131, ST1981 and ST95 (**Supplementary Table 1)**. Previously published cluster analysis showed that ST95 evolved into two separate branches, one diverged to global pandemic clone ST131 which recently evolved into ST999 and a separate divergence formed ST1981 [44]. Also, ST999, ST131, and ST95 belong to phylogeny B2, and ST1981 belongs to phylogeny D. Further, the genome size of ST999 is 4.8 Mbp and carries 187 known virulence factors (VFs). ST131 and ST1981 each have a genome size of 5.3 Mbp and carry 194 and 239 known virulence factors, respectively. ST95 has a genome size of 5.1Mbp and carries 258 known virulence factors (**Supplementary Table 1**). For comparison, the genome sizes of non-pathogenic strains XL1 blue, *E. coli* MG1655, and pathogenic pyelonephritis CFT073 are 5.2 Mbp, 4.7 Mbp, and 5.3 Mbp respectively, and they carry 142, 136, and 231 known virulence factors, respectively (**Supplementary Table 1)**. Based on fimH typing, ST999, ST131, ST1981, and ST95 are classified as *fimH20, fimH30, fimH49* and *fimH18*, respectively (**Supplementary Table 1**).

In silico O-H typing, categorized ST999, ST131, ST1981, and ST95 as *O(null):H4, O25:H4, O1:H6, O18:H7*, respectively (**Supplementary Table 1**). The ST999 genome lacks the biosynthetic wzy pathway to assemble the O-antigen. This occurred due to homologous recombination and has been previously reported [45]. Loss of the O-antigen severely impairs virulence capacity, and makes a strain more susceptible to lysozyme and complement-mediated killing [22]. In contrast, antigen types O1 and O18 which are carried by ST1981 and ST95 are known to confer resistance to serum mediated killing [46].

ST999, ST131, ST1981, and ST95 are multidrug-resistant (MDR) because of the presence of the 1,233 bp *mdf(A)* gene [47] (**Supplementary Table 1**). This gene encodes a 410 amino acid long membrane protein that functions as an MDR chemotherapeutic drugs exporter [47, 48]. ST131 additionally carries aminoglycoside modifying enzymes such as *aadA5*, which encodes aminoglycoside nucleotidyl transferase that provides resistance against streptomycin [49, 50], *aac(3)-IId* encoding aminoglycoside N(3)-acetyltransferase, which provides resistance against gentamicin, kanamycin and tobramycin [50, 51], and *aac(6’)-Ib-cr* encoding acetylating aminoglycoside-(6)-N-acetyltransferase, which provides resistance against kanamycin A/B, neomycin, gentamicin and sisomicin [50]. It also carries *tet(B)* encoding a tetracycline efflux transporter, *sul1* encoding an alternative dihydropteroate synthase that has low affinity for sulfonamides [52], *dfrA17* encoding an alternate dihydrofolate reductase, which has low affinity for trimethoprim, *blaCTX-M-15, blaTEM-1B* and *blaOXA-1* encoding extended-spectrum β-lactamases that provide resistance against various chemotherapeutic drugs, *qacE* encoding the multi-drug exporter quaternary ammonium compound-resistance protein, and *catB3* and *catA1* encoding type B/A chloramphenicol O-acetyltransferase providing resistance against chloramphenicol.

### Highly virulent ST95 resembles UTI89

Using the genome of MG1655 as the reference, the genomic maps of XL1 blue, ST999, ST131, ST1981 and ST95 were constructed (**Figure 1A**). This revealed clear genotypic differences between the UPEC strains. Most importantly, ST95 was found to closely match the canonical UPEC strain UTI89, but it carries additional virulence factors such as *traJ, traT, senB*, and *ccdb* which are absent in UTI89 (**Figure 1A, B and Supplementary Table 3)**. *traJ* and *traT* are involved in the transfer of F-plasmid [53], *ccdb* encodes topoisomerase II DNA gyrase poison that is toxic to mammalian cells [54], and *senB* encodes Shigella enterotoxin TieB protein [28, 55]. As mentioned above, ST999 is likely less virulent as it carries less virulence factors and lacks genes encoding the O-antigen (**Supplementary Table 1**). Constructing the phylogenetic trees using the pan and the core genome revealed that ST999 and ST131 have a recent common ancestry, while ST95 diverged and grouped with the canonical UPEC strains UTI89 and CFT073 (**Figure 1B**).

**Figure 1.**
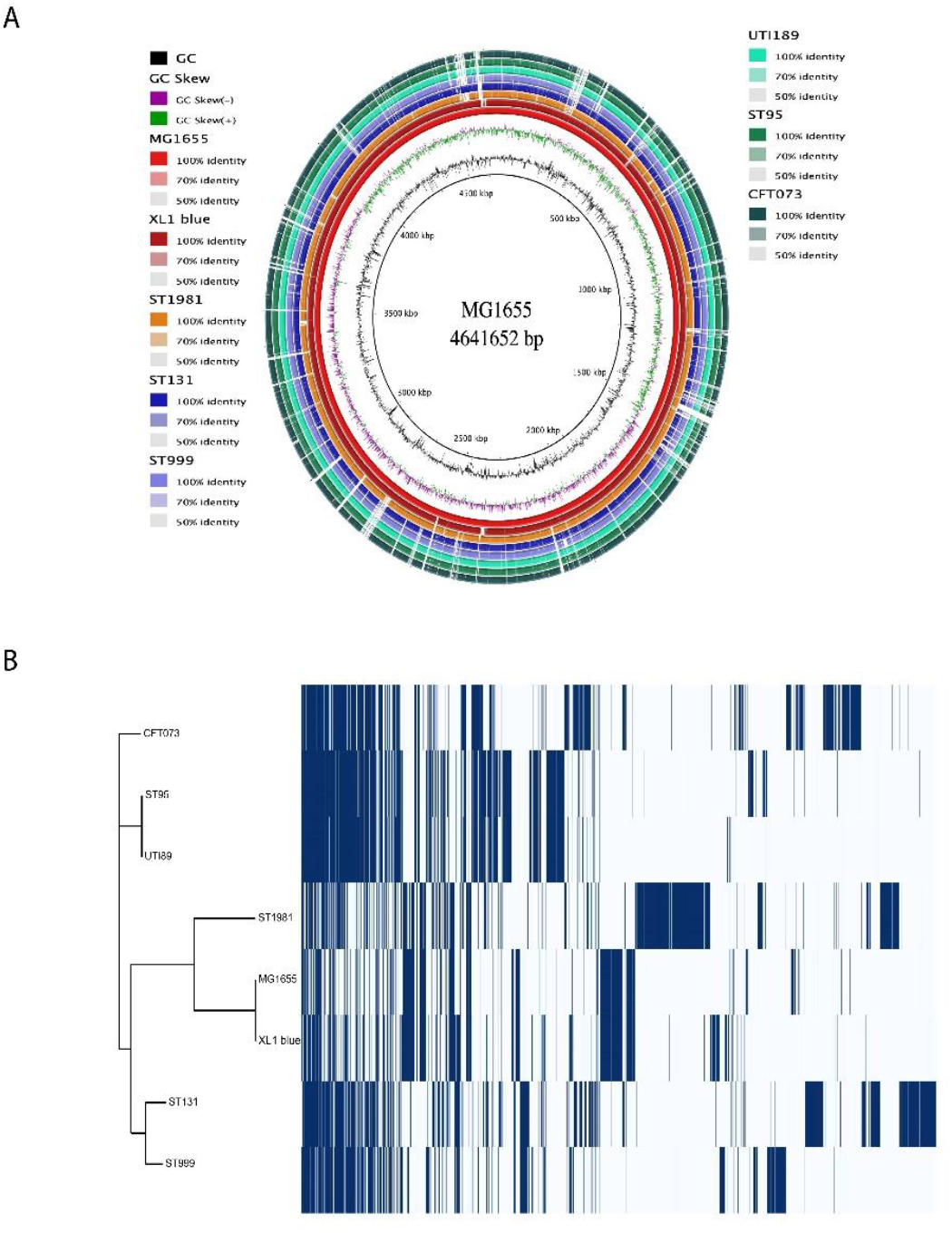
Uropathogenic strains used in this study are genetically diverse. (A) Genome maps of MG1655, XL1 blue, ST999, ST131, ST1981, and ST95 showing the absence and presence of genes. (B) Phylogenetic analysis on core (left panel) as well as pan (right panel) genome reveals that ST999 and ST131 have a recent common ancestry, while ST95 diverged from a common ancestor with UTI89 and CFT073. Blue and white lines, respectively, indicate the presence and absence of genes.

### All UPEC strains survive in macrophages

Bacterial killing by macrophages was addressed with the colony forming unit (CFU) assay. In this assay, bacteria are co-cultured with macrophages derived from peripheral blood monocytes of five different healthy donors. By subsequently killing the extracellular (i.e., not ingested) bacteria with a cocktail of antibiotics (Gentamicin, Nitrofurantoin, Imipenem, Trimethoprim, Sulfamethoxazole), the intracellular survival of bacteria within the macrophages was assessed (**Figure 2A**). All UPEC strains were sensitive to this antibiotics cocktail, as has been previously reported for multidrug resistant strains [56]. The number of bacteria was adjusted to ∼300 for infection. The counts of intramacrophagal XL1 blue, ST999, ST131, ST1981, and ST95 were 205 (± 124; *n* = 5 donors), 598 (± 301; *n* = 5), 247 (± 86; *n* = 5), 4,077 (± 2,026; *n* = 5), 186 (± 140; *n* = 5) at 4 hr post-infection (hpi), respectively. A high count of ST1981, relative to other strains suggests that ST1981 was rapidly multiplying within the macrophages. At 6 hpi, the counts of intramacrophagal XL1 blue, ST999, ST131, ST1981, and ST95 insignificantly changed to 49 (± 33; *n* = 5), 276 (± 104; *n* = 5), 212 (± 68; *n* = 5), 3,153 (± 1320; *n* = 5), and 157 (± 81.7; *n* = 5), respectively. Comparing the counts of intramacrophagal ST999, ST131, ST1981, and ST95 at 4 and 6 hpi, suggests that the macrophages were better able to clear XL1 blue than the UPEC strains.

**Figure 2.**
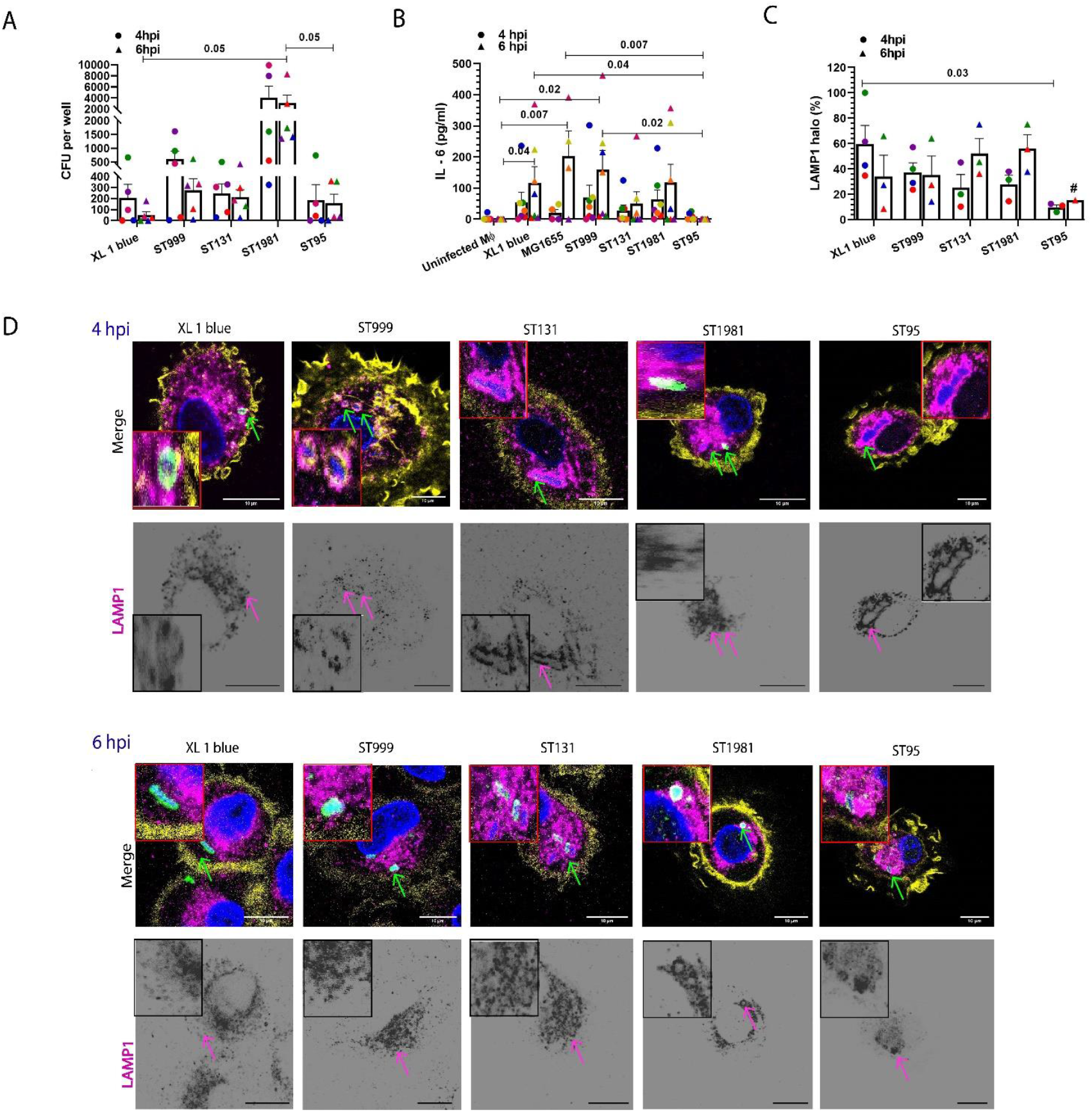
Uropathogenic strains ST999, ST131 and ST1981 trigger IL-6 production and survive within macrophages. (A) Colony forming unit (CFU) assay showing intracellular survival of UPEC strains ST999, ST131, ST1981 and ST95 and XL1 blue at 4 and 6 hr post-infection (hpi). Each color represents an individual donor. (n = 5 donors, two-tailed unpaired t-test). (B) IL-6 secretion by the macrophages. (n = 4-7 donors, two-tailed unpaired t-test). (C) Percentage of phagosomes carrying *E. coli* with positive immunostaining for the late endosomal/lysosomal marker LAMP1. (n = 3-4 donors, two-tailed unpaired t-test). (D) Representative confocal microscopy images with immunostaining for LAMP1 (magenta in merge) and LPS (green). Yellow: phalloidin. Blue: DAPI. Scale bars, 10 μm. Note that bacterial DNA is also stained with DAPI. # indicate no intact macrophages were observed in two experiments.

It has been reported that within ∼2 hr following the ingestion of bacteria by macrophages, the bacteria-containing vacuoles fuse with LAMP1 positive late endosomal and lysosomal compartments [17]. Thereby, these vacuoles acquire bactericidal properties such as lytic enzymes and low pH. We therefore assessed LAMP1 recruitment to the bacteria-containing vacuoles (**Figure 2C, D**). At 4 hpi, recruitment of LAMP1 to XL1 blue, ST999, ST131, ST1981, and ST95-containing vacuoles was 60% (± 14; *n* = 4 donors), 38% (± 7.3; *n* = 4), 25% (± 10; *n* = 3), 28% (± 7; *n* = 3), and 10% (± 2; *n* = 3), respectively. LAMP1 recruitment did not change significantly at 6 hpi. Thus, for all UPEC strains, 10 – 38% of all vacuoles acquired the late endosomal/lysosomal marker LAMP1, whereas 60% of XL1 blue carrying vacuoles contained LAMP1.

### ST95 does not induce inflammatory cytokine production by macrophages

Immunogenicity of ST999, ST131, ST1981, and ST95 was assessed by measuring the levels of secreted interleukin 6 (IL-6) (**Figure 2B**) and tumor necrosis factor alpha (TNF-α) (**Supplementary Figure 1B**). Basal levels of IL-6 detected in uninfected macrophages were 3 pg/ml (± 3; *n* = 7 donors). At 6 hpi, IL-6 levels significantly increased to 117 pg/ml (± 52; *n* = 7), 202 pg/ml (± 82; *n* = 4), and 159 pg/ml (± 63; *n* = 7) in XL1 blue, MG1655 and ST999 infected macrophages, respectively. Likely due to large donor variation, no significant increases in IL-6 production occurred in ST131, ST1981 and ST95 infected macrophages. Basal levels of TNF-α detected in uninfected macrophages were 12 pg/ml (± 8; *n* = 6) at 4 hpi (**Supplementary Figure 1B**). A significant increase in TNF-α levels to 24 pg/ml (± 8; *n* = 6) occurred in ST131 infected macrophages. Also, at 6 hpi, significant increases to 59 pg/ml (± 23; *n* = 4) and 19 pg/ml (± 7; *n* = 6) occurred in MG1655 and ST131 infected macrophages. The levels of secreted IL-6 and TNF-α were only 0.0 pg/ml (± 0.0; *n* = 7) and 3.4 pg/ml (± 3.4; *n* = 6) in ST95 infected macrophages at 6 hpi, and, in contrast to the other *E. coli* strains, these levels did not consistently increase between 4 and 6 hpi, indicating that ST95 was non-immunogenic under these conditions (**Figure 2C and Supplementary Figure 1B**). These data show that XL1 blue, MG1655, ST999, and ST131 induced inflammatory cytokine production by the macrophages, whereas ST95 did not induce secretion of these cytokines.

### ST95 produce haemolysin and induced macrophage death

The whole-genome sequencing revealed that ST95 carries hemolysin genes *hlyABCD* while ST999, ST131, and ST1981 lack these genes (**Table 1**). Haemolysin induces nod-like receptor family pyrin domain containing 3 (NLRP3) dependent or independent cell death of macrophages, depending on its concentration [57]. In the CFU assay, the counts of intramacrophagal ST95 were lower than other UPEC strains and, in contrast to XL1 blue, did not seem to change over time (**Figure 2A**). Moreover, we found that ST95 did not elicit cytokine responses from the cultured macrophages (**Figure 2C**). Importantly, at 4 hpi, we found in the first donor that the recruitment of LAMP1 to ST95-containing vacuoles was only 15.0% (*n* = 1), and for the remaining two donors we did not find intact macrophages in our microscopy experiments (**Figure 2B**). These findings raised the possibility that ST95 was triggering macrophages death.

**Table 1.**
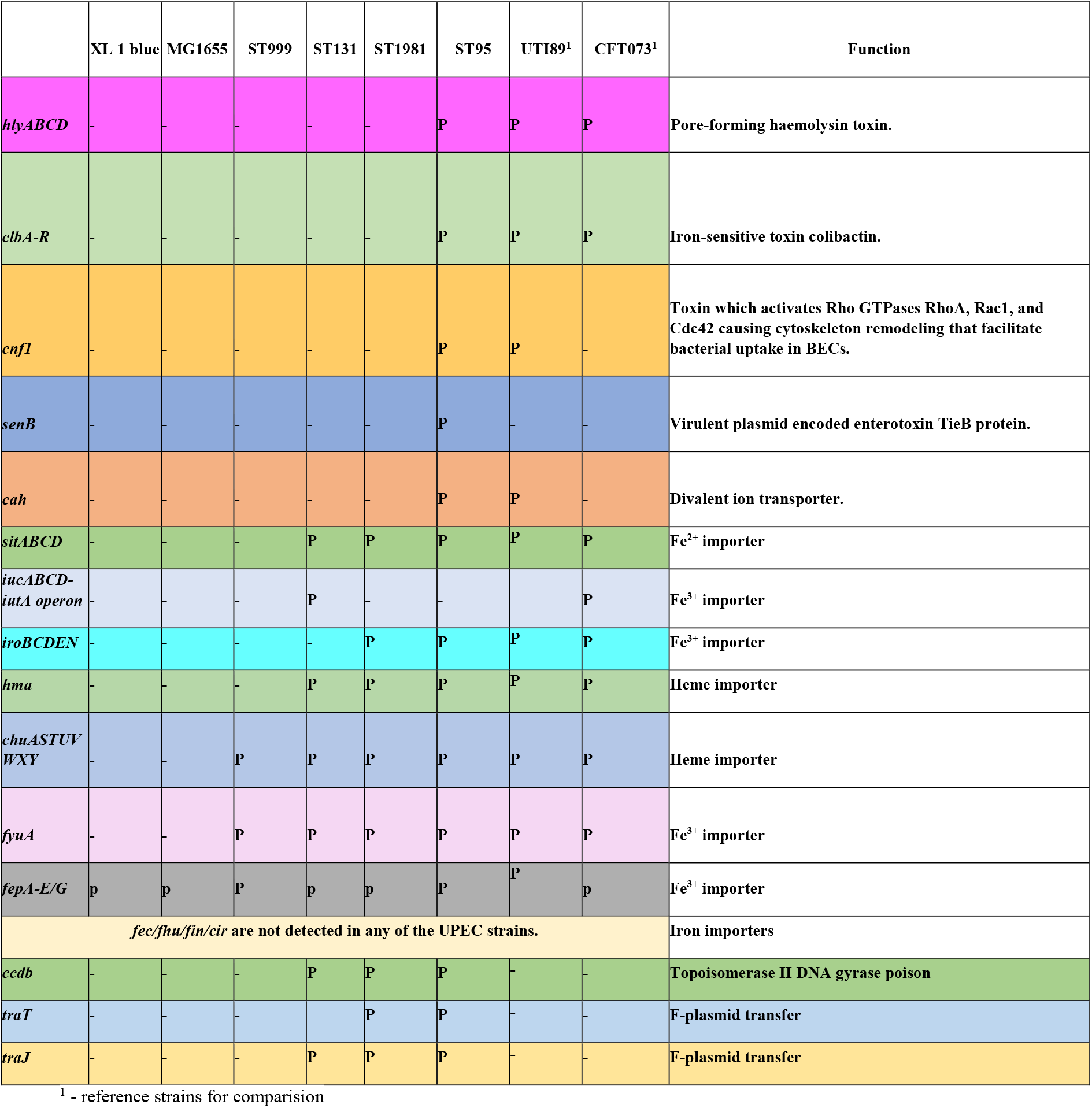
Iron acquisition genes and toxins found in non-pathogenic *E. coli* and UPEC strains of this study.

To confirm that ST95 is inducing macrophages cell death, we next measured propidium iodide intensity in ST95 infected macrophages at 1 hpi (**Figure 3A**). A significantly higher PI positive population indicating dead macrophages was observed in ST95 infected macrophages, relative to the other *E. coli* strains (**Figure 3A)**. We also assessed hemolytic activity of the UPEC strains. For this, lysates of the 1 hpi macrophages were cultured on a blood agar plate (**Figure 3B**). The discoloration of the red blood cells at the sites of culturing lysates of ST95 infected macrophages indicates hemolytic activity. In contrast, no discolorations were observed at the sites of culturing lysates of XL1 blue, MG1655, ST999, ST131, and ST1981 infected macrophages.

**Figure 3.**
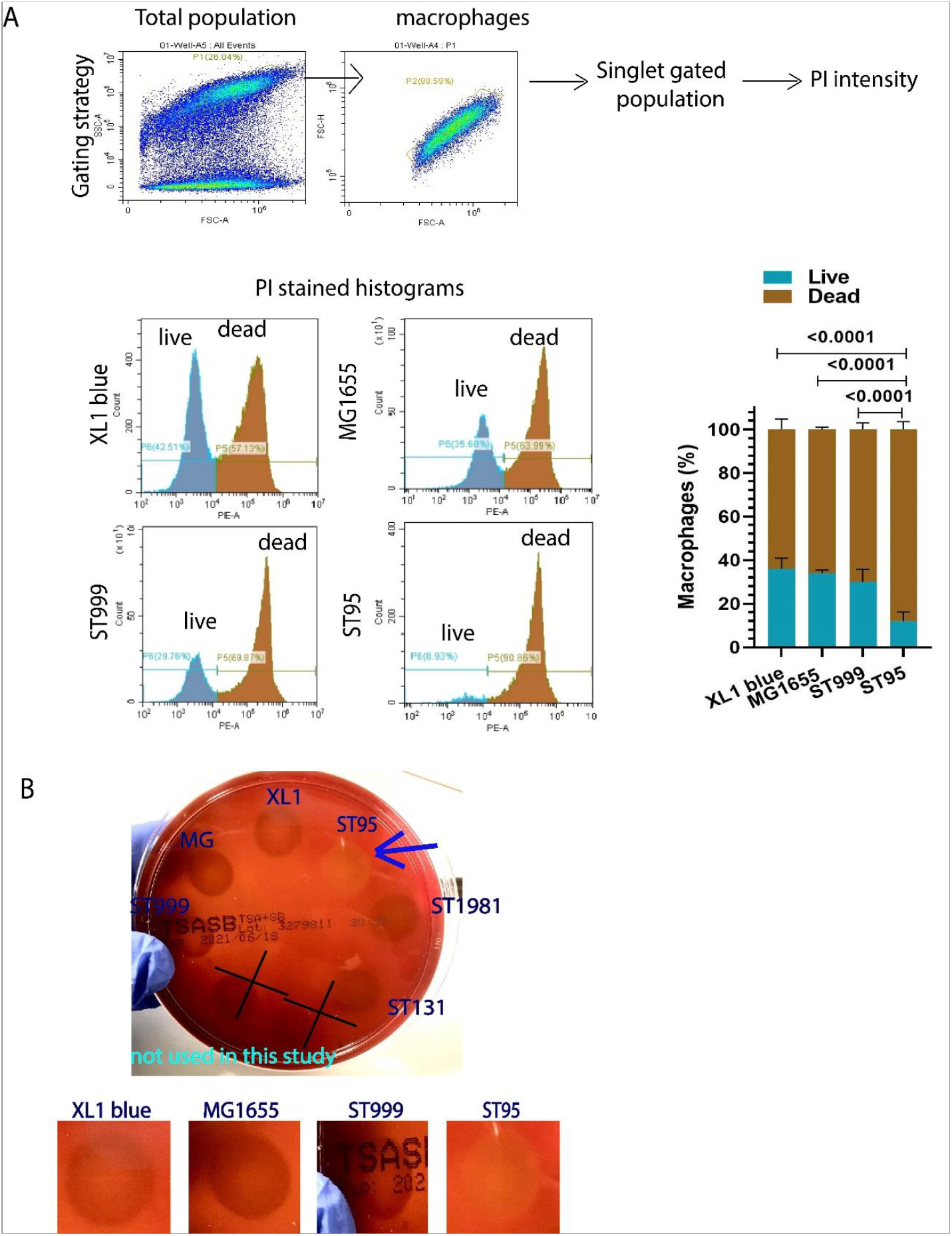
Lytic activity and induction of macrophage death by ST95. (A) Flow cytometry gating strategy and representative histograms. Cells were selected on side and forward scatter (SSC, FSC) area (A) and height (H). Macrophages were stained with the life/death marker propidium iodide (PI) showing the percentage of live and dead macrophage population. Infection was carried out at the ratio of 10 macrophages per bacterium. (n = 4 one way donors, one-way ANOVA with Tukey’s multiple comparisons test). (B) Representative blood agar plate with colonies of the UPEC strains isolated from 1-hour post infected macrophages. Arrow: hemolytic activity of ST95.

### ST999, ST131 and ST1981 reside in double membrane vacuoles in macrophages

UPEC has been reported to induce a specialized form of autophagy, called xenophagy, the process by which cells direct the autophagy machinery against pathogens [58, 59]. Therefore, we assessed whether the UPEC strains activated xenophagy in infected macrophages by electron microscopy. The presence of bacteria containing double membranes vacuoles indicates activation of xenophagy [60, 61]. The percentages of double membrane vacuoles in ST999, ST131, and ST1981 infected macrophages were 47%, 51%, and 46%, respectively, whereas only 0% of XL1 blue and 22% of MG1655 resided in double membrane vacuoles (**Figure 4A, and supplementary Figure 1, 3, and 4**).

**Figure 4.**
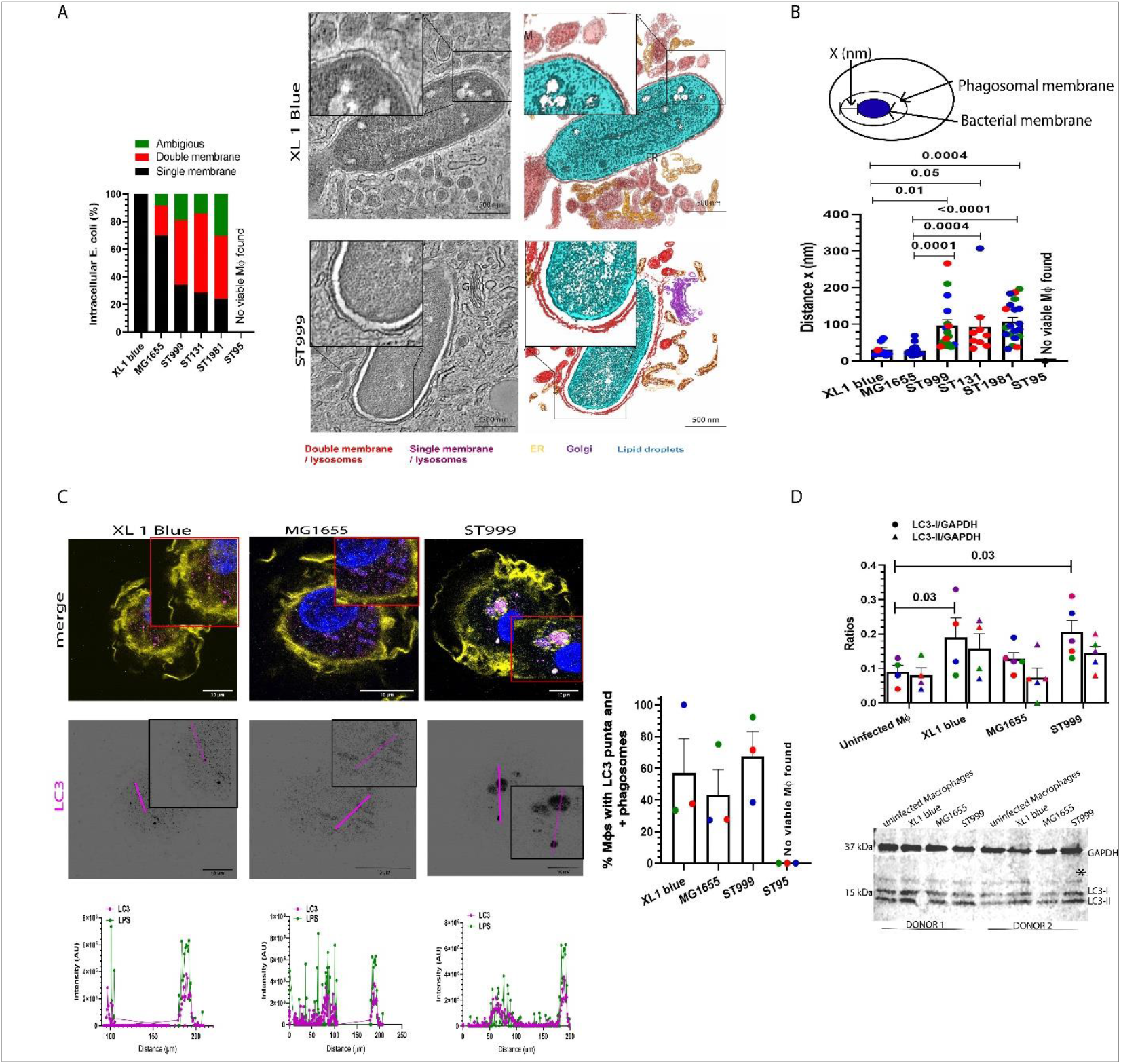
UPEC strains reside in double-membrane autophagosomes, whereas non-pathogenic *E. coli* reside in single-membrane LC3-associated phagosomes. (A) Left: Percentage of *E. coli* carrying phagosomes with double and single membranes (7 - 60 phagosomes pooled from n = 2-3 donors). Right: Representative tomogram with single (upper) and double (lower) membrane phagosomes containing XL1 blue and ST999, respectively. (B) Distance between the outer membrane of the bacteria and the phagosomal membrane. Each color represents an individual donor. (n = 2-3 donors, two-tailed unpaired t-test). (C) Left: representative confocal images of macrophages infected with *E. coli* for 1 hr and immuno stained for LC3 at 4 h post infection. LC3: magenta in merge. LPS: green. Yellow: phalloidin. Blue: DAPI. Scale bars, 10 μm. Graphs: fluorescence intensity cross sections as indicated. Right: percentages of macrophages with LC3 positive punta/compartments at 4-hour post-infection. (D) Representative western blot of *E. coli* infected macrophages stained for LC3 and GAPDH (loading control). LC3-I: non conjugated form of LC3. LC3-II: lipid conjugated form of LC3. Graph: Densitometry quantification of 3 experiments showing LC3-I and LC3-II levels in infected macrophages. *unspecific band.

From the electron microscopy micrographs, we also measured the distance between the outer membrane of the bacteria and the (inner or limiting) membrane of the vacuoles at 1 hpi (**Figure 4B**), because the presence of a large space around bacteria in vacuoles is suggestive of proliferating bacteria [62-64]. The distances between the phagosomal membrane and the bacterial outer membranes were 96 nm (± 16.6; *n* = 32), 115 nm (± 46; *n* = 10), and 109 nm (± 15; *n* = 35) for ST999, ST131 and ST1981, respectively. These distances were significantly lower for XL1 blue and MG1655: 29 nm (± 5; *n* = 18) and 27 nm (± 3; *n* = 53), respectively (**Figure 4B**). This suggests that ST999, ST131 and ST1981 might multiply within the vacuoles, in line with our findings from the CFU assay (**Figure 2A and Supplementary Figure 1**), and the occasional occurrence of multiple bacteria within the vacuoles observed by electron microscopy (**Supplementary Figure 3)** and fluorescence microscopy **(Supplementary Video 14**).

In the next set of experiments, we performed side-by-side comparisons between ST999 and ST95. We focused on these UPEC strains, because these were the most different in terms of number of virulence factors (**Table 1 and Supplementary Table 1**). To confirm that xenophagy was activated in ST999 infected macrophages, we assessed the presence of LC3 punta/positive vacuoles by immunofluorescence microscopy. Surprisingly, although we observed clear LC3 staining in ST999 infected macrophages, this did not significantly differ from the non-pathogenic strains XL1 blue and MG1655. The percentages of cells with LC3 punta/positive vacuoles were 57% (± 22; *n* = 3 donors), 43% (± 16; *n* = 3), and 67% (± 16; *n* = 3) in XL1 blue, MG1655, and ST999 infected macrophages, respectively (**Figure 4C**). Western blot also revealed no significant differences between the *E. coli* strains, and XL1 blue and ST999 both increased the levels of non-lipidated LC3-I (**Figure 4D**). Collectively with LC3 staining and that XL1 blue and MG1655 reside in single membrane vacuoles while ST999 resides in double membrane vacuoles (**Figure 4A**), suggests that XL1 blue and MG1566 were internalized via LC3 associated phagocytosis (LAP) while ST999 induces xenophagy [61]. In LAP, LC3 is directly recruited to the limiting membrane of the vacuoles in LAP, resulting in efficient phagosome-lysosome fusion and degradation of the cargo as has been reported previously for other bacterial pathogens [65, 66].

### ST999 and ST95 survive in epithelial cells

Next, we determined whether ST95 also induced cell death of BECs. We first confirmed that both ST999 and ST95 expressed fimH which is needed for infection of BECs by interaction with uroplakin Ia [67] (**Supplementary Figure 2A, B)**. Next, we carried out CFU assays to determine the survival of bacteria within the BECs. Infections were carried out on 100% confluent BECs at the ratio of 1,500 BECs per bacterium. At 4 hpi, the counts of intracellular XL1 blue, MG1655, ST999 and ST95 were similar (**Figure 5A)**. Importantly, the LDH leakage assay showed that ST95 did not cause BECs death, even at the ratio of 10 BECs per bacterium (i.e., the same as used for the macrophages) (**Figure 5B)**. We assessed whether the UPEC strains displayed hemolytic activity using the blood agar plate assay (**Figure 5C**). Similar to the results with macrophages, no discoloration of the red blood cells was observed at the sites of the culturing lysates of XL1 blue, MG1655, ST999, ST1981 and ST131 infected BECs. However, we observed discoloring at the site of culturing ST95. Thus, although ST95 exerts hemolytic activity, this does not seem to suffice for killing of the BECs.

**Figure 5.**
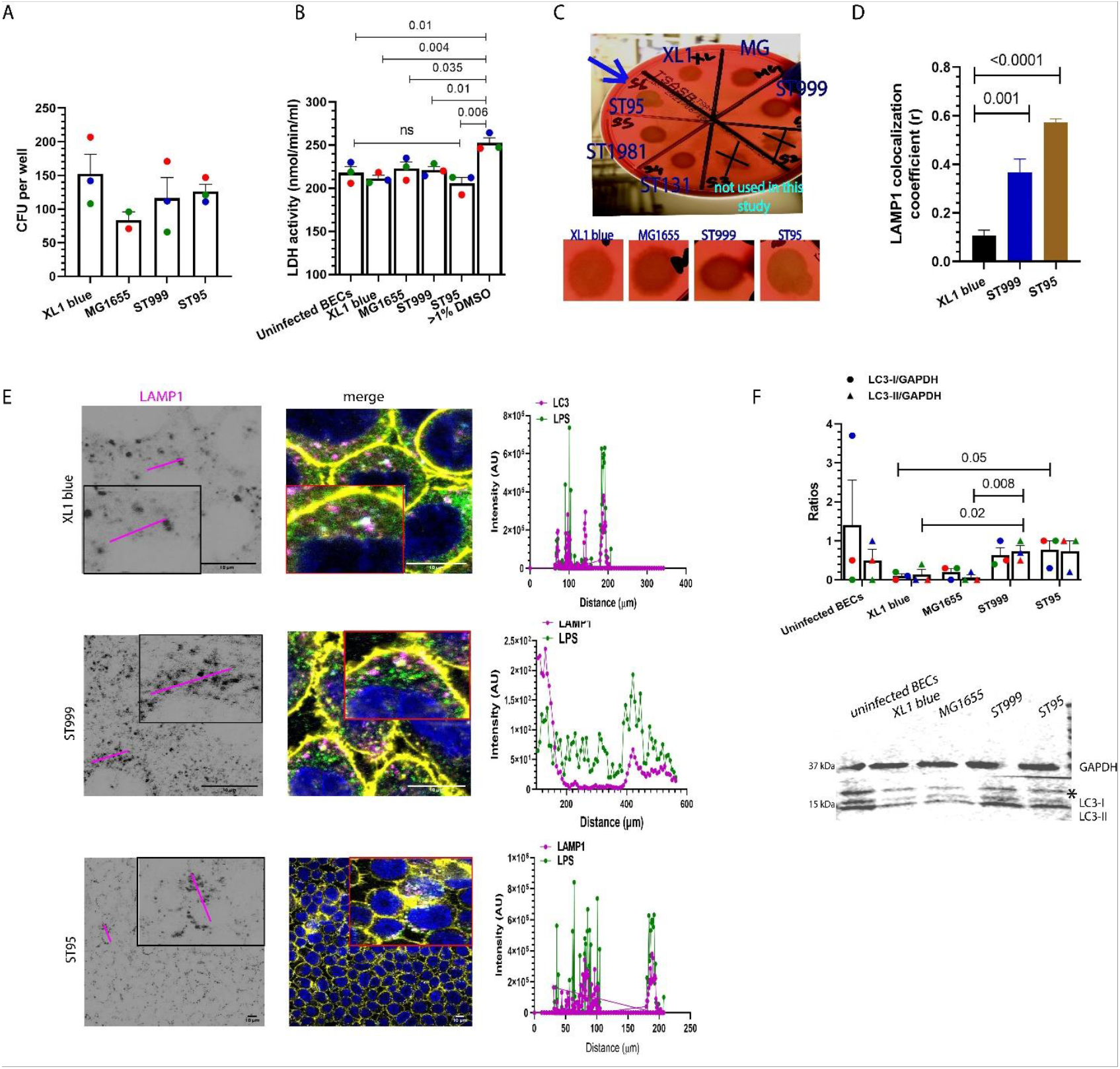
ST95 does not induce cell death in BECs. (A) Colony forming unit (CFU) assay showing intracellular survival of *E. coli* strains ST999, ST95, MG1655 and XL1 blue at 4 hr post-infection (hpi). Each color represents an individual donor. (n = 3 experiments). (B) BEC death assessed by leakage of lactate dehydrogenase (LDH). DMSO: positive control. 10 BECs per *E. coli* were used. Each color represents a individual donor. (n = 3 experiments, two-tailed unpaired t-test). (C) Blood agar plate with colonies of the UPEC strains isolated from 1-hour post infected BECs. Arrow: hemolytic activity of ST95. (D) Percentage of *E. coli* co-localized with late endosomal/lysosomal marker LAMP1 at 4 hr post-infection. (n = 372 - 1865 cells pooled from 3 experiments, two-tailed unpaired t-test). (E) Representative confocal microscopy images with immunostaining for LAMP1 (magenta in merge) and LPS (green). Yellow: phalloidin. Cyan: DAPI. Scale bars, 10 μm. Graphs: fluorescence intensity cross-sections as indicated. (F) Densitometry quantification of 3 experiments showing LC3-I and LC3-II levels in infected BECs. Lower panel: Representative western blot of *E. coli* infected BECs stained for LC3 and GAPDH. LC3-I: non conjugated form of LC3. LC3-II: lipid conjugated form of LC3. Each color represents an individual donor. *unspecific band.

Within the BECs, both ST999 and ST95 resided in more LAMP1 positive compartments than XL1 blue at 4hpi (**Figure 5D, E**). The Pearson’s co-localization coefficients of intracellular XL1 blue, ST999, and ST95 with LAMP1 were 0.11 (± 0.02; *n* = 372 cells), 0.37 (± 0.05; *n* = 1,689), and 0.57 (± 0.02; *n* = 1,859), respectively (**Figure 5D**). Moreover, we found that ST999 and ST95 increased lipidated LC3-II levels (**Figure 5F**), as has been previously reported for UPEC infected epithelial cells [68].

### Iron supplementation differentially affected growth of ST999 and ST95

In order to survive in the urinal tract, UPEC has to acquire sufficient iron. To accomplish this, UPEC strains carry redundant iron acquisition systems [23]. These iron acquisition systems include haem transporters (*chu, hma*), ferrous ions (Fe^2+^) importer (*sit*), Enterobactin siderophore system (*fep*), Salmochelin siderophore system (*iro*), Aerobactin siderophore system (*iuc, iut*) and Yersiniabactin siderophore system (*fyu*). All siderophores import ferric ions (Fe^3+^). Bioinformatics analysis revealed that different combinations of these iron acquisition genes are present in ST999, ST131, ST1981, and ST95 (**Table 1**). ST999 carries *chu, fep*, and *fyuA* genes to import haem and Fe^3+^, but lacks Fe^2+^ ion (*sitABCD*) importer. ST131 carries *chu, hma, sit, fep, iuc* and *fyuA* genes to import heme, Fe^2+^ and Fe^3+^. Both ST1981 and ST95 carry *chu*, and an additional *hma to import heme. They both carry sit, fep, iro* and *fyuA* genes to import Fe^2+^ and Fe^3+^ (**Table 1)**.

Since the UPEC strains carry different combination of iron acquisition genes, we next assessed how iron affected the growth of the ST999 and ST95 strains. We again selected ST999 and ST95 for assessing their iron tolerance, because these strains carry the minimum and maximum number of iron acquisition genes, respectively, and they differently affected the macrophages. We first measured the growth rate in mammalian cell culture medium: RPMI-1640 media containing 10% FBS, which has an estimated 5.6 ± 1.3 μM iron concentration [69] (**Figure 6A**). Both ST999 and ST95 showed similar growth rates, whereas growth of XL1 blue was significantly slower. To assess how iron affected their growth, ST999 and ST95 were grown in minimal M9 medium (no iron added, but might contain contaminations of iron), M9 medium supplemented with 0.1 mM FeCl^3^ and M9 medium with 1 µM of the iron chelator deferoxamine (**Figure 6)**. The growth rate of ST999 was accelerated in the presence of Fe^3+^, but was significantly slowed upon iron chelation. In contrast, the growth rate of ST95 was significantly reduced in the presence of Fe^3+^ while no significant change occurred upon iron chelation. Growth of the control strain XL1 blue was slowed both with iron supplementation and chelation. This shows that ST999 grows better in the presence of high Fe^3+^ concentrations, whereas ST95 grows better at low Fe^3+^ concentrations.

**Figure 6.**
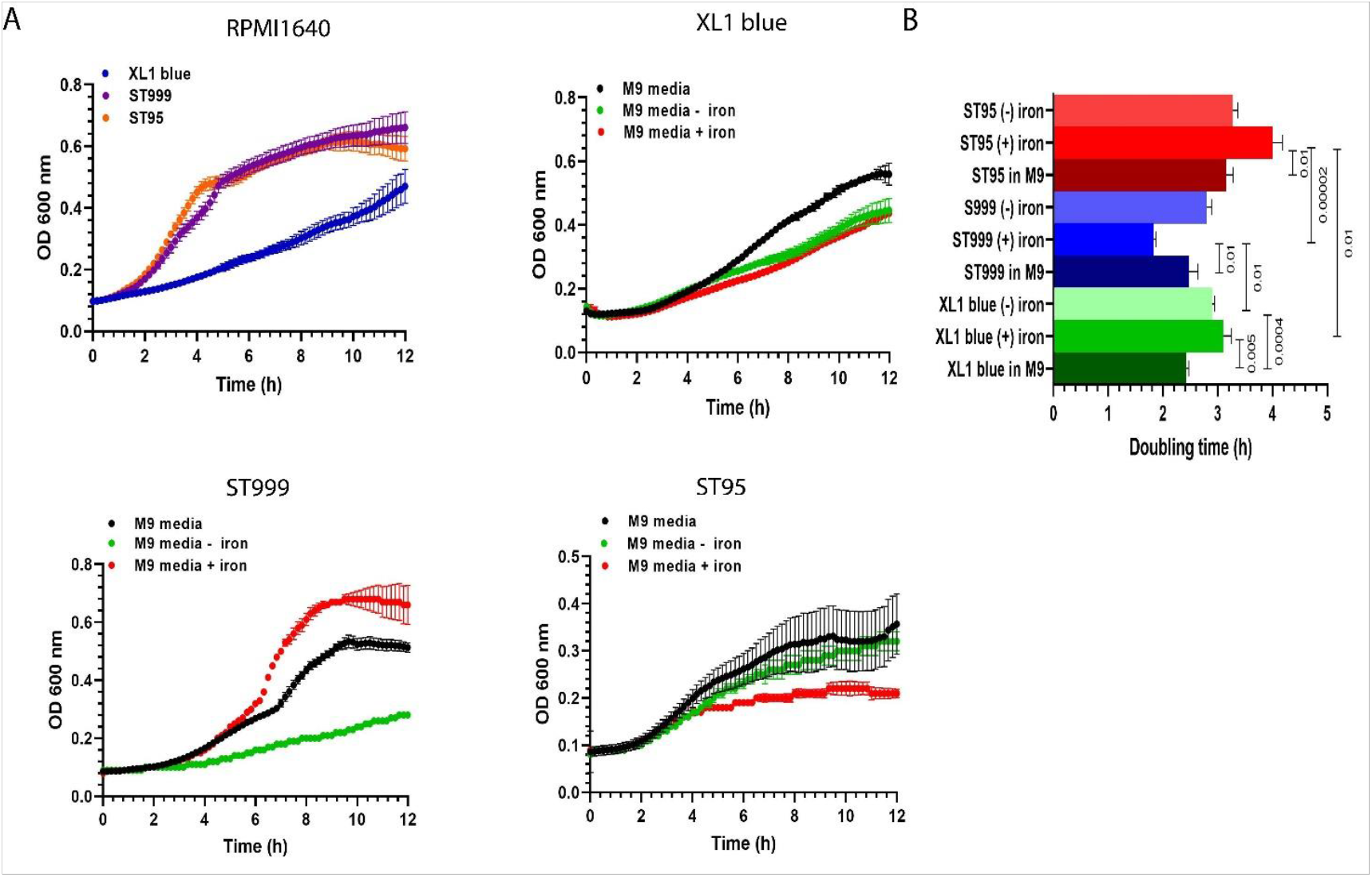
Differential iron requirement for growth by UPEC. (A) Growth curves of XL1 blue, ST999 and ST95 in (i) RPMI medium supplemented with 10% FBS, M9 medium (black curves in subpanels ii – iv), M9 medium with 0.1 mM FeCl3 (red curves), and M9 medium with 1 μM deferoxamine (green curves). Average ± SEM. (B) Average doubling times in exponential growth phase. (n = 3-4 experiments, two-tailed unpaired t-test).

We next assessed whether iron affected the expression of virulence factors in ST999 and ST95 by RT-qPCR. Normalized to the expression levels in M9 media, the fold expression of *chuA* and *oxyR* insignificantly increased to 8.8 (± 7.9; *n* = 4) and 3.0 (± 2.5; *n* = 4) in ST999 grown in M9 media supplemented with FeCl^3^, respectively (**Figure 7A**), and changed to 1.5 (± 0.8; *n* = 4) and 0.5 (± 0.2; *n* = 4) in M9 media supplemented with deferoxamine, respectively. Iron induced opposing effects in the expression of *chuA* in ST95. Normalized to ST95 grown in M9 media, fold expression of *chuA* non-significantly decreased to 0.3 (± 0.1; *n* = 4) and 0.3 (± 0.04; *n* = 4) in ST95 grown in M9 media supplemented with FeCl^3^ and deferoxamine, respectively. The fold expression of *oxyR* in ST95 showed a similar pattern as observed for ST999. However, iron availability caused mixed patterns in the expression of other the virulence factors *clbN, senB, hma, iroN* and *fliC* (**Figure 7B**). Importantly, the fold expression of *clbN* significantly increased to 19.5 (± 11.5; *n* = 5) in ST95 grown in M9 media supplemented with deferoxamine and remained low at 0.9 (± 0.01; *n* = 3) in M9 media supplemented with FeCl^3^ (**Figure 7B**). Similarly, the fold expression of *senB* also increased non-significantly to 1.9 (± 0.4; *n* = 5) in M9 media supplemented with deferoxamine while was low at 1.3 (± 0.4; *n* = 4) in M9 media supplemented with FeCl^3^ (**Figure 7B**).

**Figure 7.**
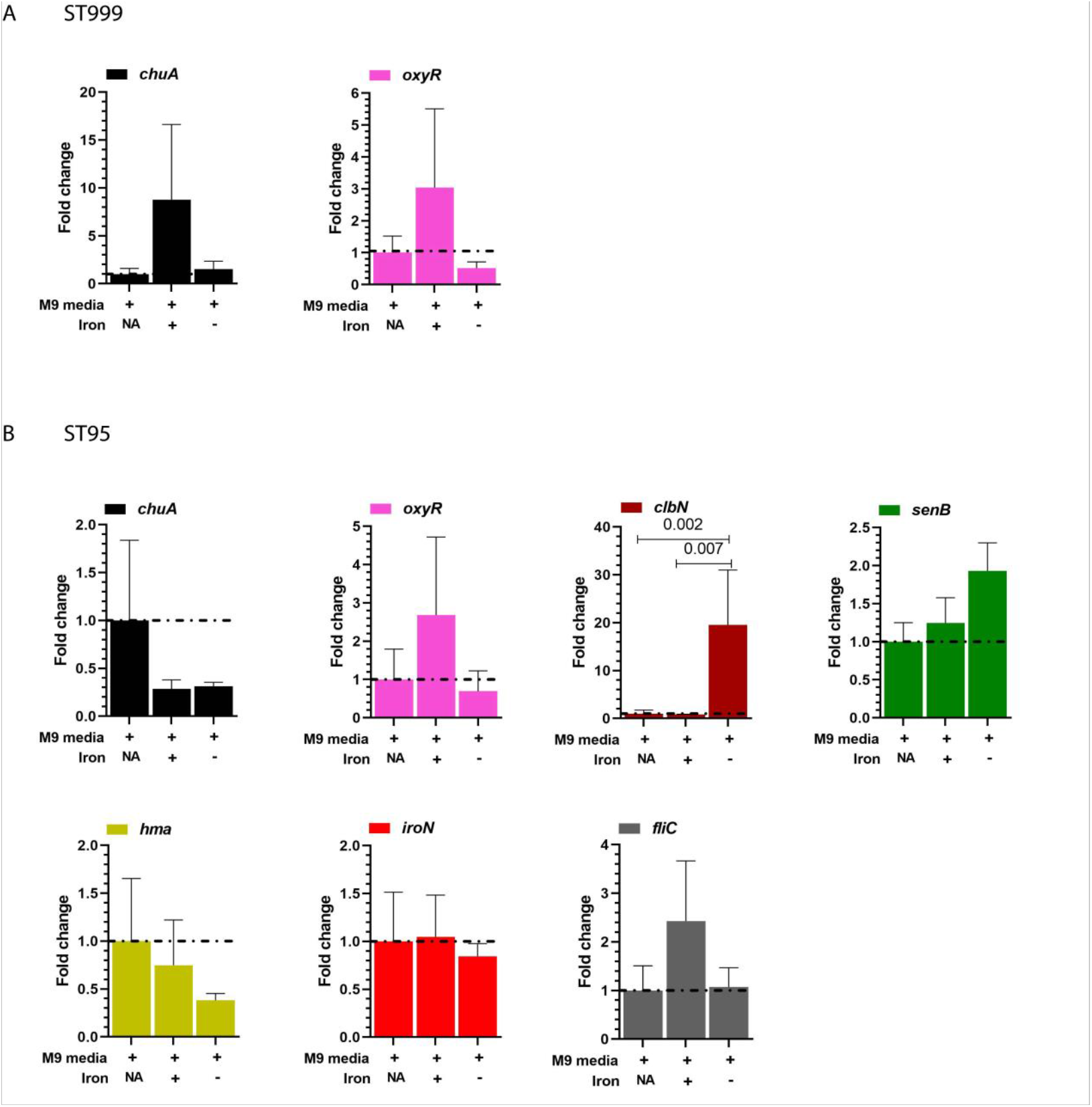
Differential expression of virulence factors by UPEC. (A) Expression of *chuA* and *oxyR* by ST999 in M9 medium, M9 medium with 0.1 mM FeCl3, and M9 medium with 1 μM deferoxamine indicated by plus and minus symbol, respectively. NA: not added. (n = 3-4 independent experiments, two-way ANOVA with Tukey’s multiple comparisons test). (B) Expression of *chuA, oxyR, clbN, senB, hma, iroN*, and *fliC* by ST95 in the presence of FeCl3 and deferoxamine. (n = 3-5, two-way ANOVA with Tukey’s multiple comparisons test). Data are relative to 16s rRNA and normalized to the M9 condition.

One way how UPEC can acquire iron, alternative to the expression of high-affinity uptake systems, is by surviving in late endosomes and lysosomes, as these compartments are the major pool of labile iron in mammalian cells [70]. In order to determine whether ST999 would locate to iron rich vacuoles, we carried out time-lapse fluorescence microscopy experiments of macrophages labeled with the iron probe Mem-RhoNox [42]. This probe become fluorescent upon deoxygenation in the presence of oxygen. The bacteria were labeled with AlexaFluor (AF) 633 C5-maleimide (**Figure 8, Supplementary Videos 1 - 13**). ST999 positive compartments stained highly positive for Mem-RhoNox (**Figure 8, Supplementary videos 4 - 8)**. In contrast, we did not observe clear Mem-RhoNox staining for ST95 and in many cases observed macrophage death (apparent from macrophage blebbing and swelling) (**Figure 8, Supplementary videos 9 - 13**). XL1 blue also located in iron-rich vacuoles (**Figure 8, Supplementary video 1 - 3)**. Thus, our data show that ST999 survives as an intracellular pathogen in iron-rich double-membrane vacuoles, whereas ST95 induces macrophage death and is better adapted to survive in the low-iron urinary tract.

**Figure 8.**
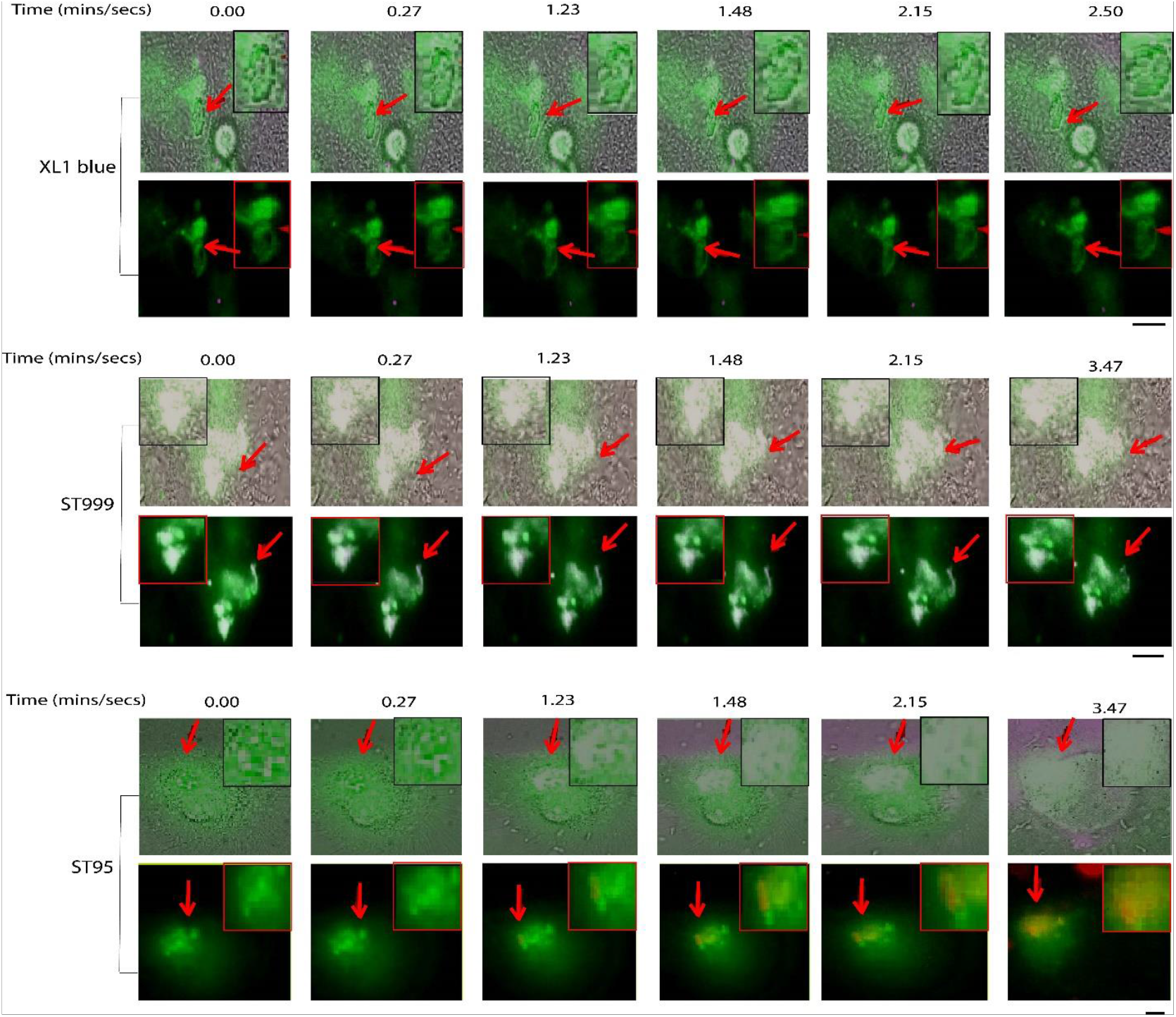
XL1 blue and ST999 reside in iron rich vacuoles, and ST95 induces macrophage death. Time-lapse microscopy of macrophages labeled with the iron probe Mem-RhoNox (green) that becomes more fluorescent in the presence of oxygen. Note that ST999 is stained positive for Mem-RhoNox and ST95 induced cell death as apparent from macrophage swelling and blebbing. Red arrows: bacteria. Scale bars, 10 μm. Also see Supplementary movies 1 - 13.

### *chuA* is expressed in macrophages, but not in BECs

Next, we determined the expression of virulence factors by ST999 and ST95 in macrophages and BECs. In macrophages, *chuA* was expressed by both ST999 and ST95 (**Figure 9A)**. However, we did not detect the *chuA* transcript in ST999 and ST95 infected BECs. In contrast, the expression of *oxyR* and the other ST95 specific virulence factors *iroN, fliC, senB* and *hma* did not significantly differ between macrophages and BECs (**Figure 9A)**. *ClbN* was undetected in ST95 infected macrophages as well as BECs.

**Figure 9.**
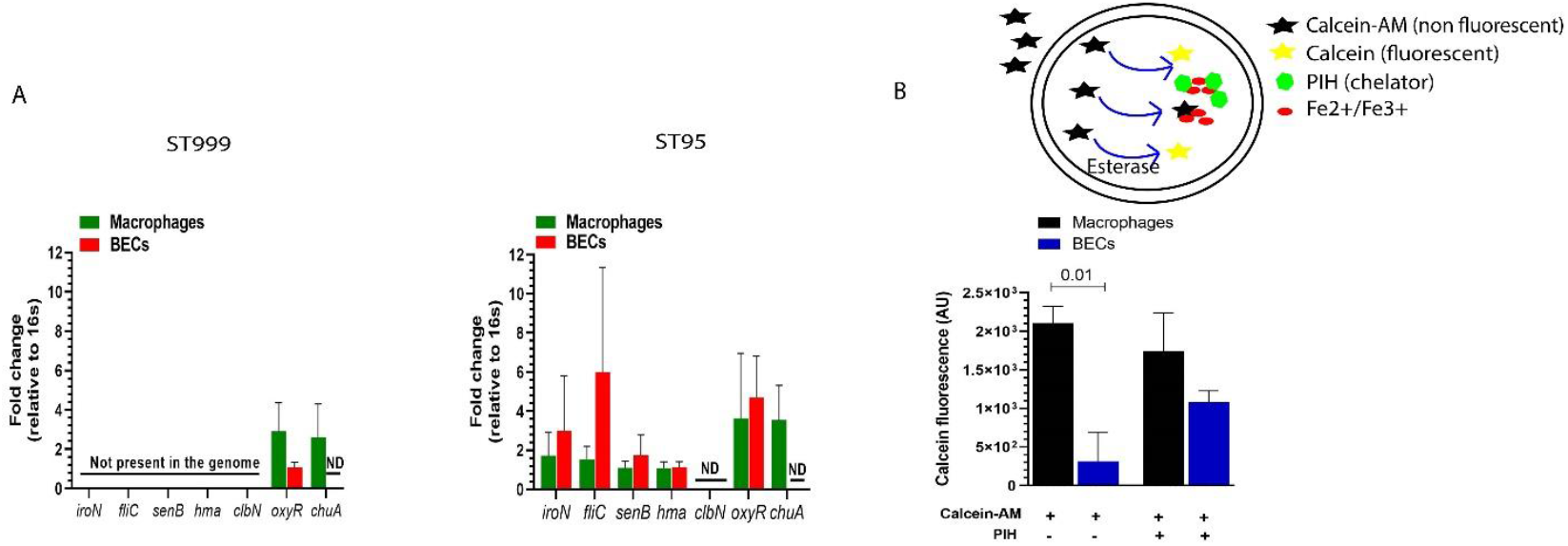
*chuA* is expressed by ST999 and ST95 following infection of macrophages but not BECs. (A) Expression of *iroN, fliC, senB, hma, clbN, oxyR* and *chuA* in infected macrophages and BECs by ST999 (right) and ST95 (left) relative to 16s RNA by RT-qPCR at 1 hr post-infection. ND: not detected. (B) Labile iron pool in macrophages and BECs measured by calcein-AM. Lower fluorescence indicates more labile iron. PIH: pyridoxal isonicotinoyl hydrazone iron chelator negative control. (n = 3 experiments, two-tailed unpaired t-test.).

Because we observed that *chuA* expression in ST999 and ST95 depended on iron availability (**Figure 7**), we next measured the labile iron pool in macrophages and BECs using calcein-AM (**Figure 9B**). Calcein-AM is membrane permeable and accumulates in the cytoplasm where the action of esterases converts calcein-AM to membrane impermeable fluorescent calcein [71]. Intracellular iron quenches the fluorescence of calcein and the fluorescence signal is thus inversely proportional to the concentration of iron [71]. Calcein fluorescence detected in macrophages and BECs was 2,108 (arbitrary units; ± 213; *n* = 3 donors), and 313 (± 381; *n* = 3 independent repeats), respectively. The significantly lower fluorescence of calcein in BECs, relative to macrophages, indicates a higher labile iron pool in BECs than macrophages. To confirm this, the labile iron was chelated with pyridoxal isonicotinoyl hydrazone (PIH) for 10 mins, before incubation with calcein-AM. After iron chelation by PIH, the calcein fluorescence increased to 1,085 (± 143; *n* = 3) in BECs thus confirming that the labile iron pool in BECs is chelatable (**Figure 9B)**. These findings indicate that BECs have a larger pool of labile iron compared to macrophages, and therefore contributed to different expression of *chuA* in BECs and macrophages.

## Discussion

In order to survive in the nutrient-limited conditions of the urinary tract, UPEC needs to acquire sufficient iron, avoid flushing in the urine by invading BECs, and evade clearance by immune cells. Overall, UPEC achieves this by invading mammalian cells, where it is shielded from surveilling macrophages and gains access to intracellular iron and other nutrients [11, 12]. However, we found differences between UPEC strains and between macrophages and BECs.

Intracellular iron is present as free Fe^2+^ and Fe^3+^ cations, called the labile iron pool, and in a protein bound form by complexing with for example porphyrin or haem co-factors [70]. The lumen of most organellar compartments, including phagosomes and lysosomes, are oxidative due to the action of enzymes involved in Fenton and Haber Weiss chemistry [17]. Hence, the oxidized Fe^3+^ form predominates in the (auto)phagosomes where intracellular UPEC resides, whereas the Fe^2+^ form predominates in the reductive cytosol [70]. The estimated concentration of labile iron in endo/lysosomes of rat liver endothelial cells has been determined as 16.0 ± 4.0 μM, whereas it is 7.3 ± 2.6 μM in the cytosol [70]. We found that ST999 carries systems for importing the oxidized Fe^3+^ form (*fep* and *fyuA*) and haem-bound form of iron (*chuA*), but lacks known importers for the reduced Fe^2+^ form (*sitABCD*). Therefore, the Fe^3+^ rich phagolysosome in BECs might be the optimal environment for the growth of ST999. This is consistent with the significantly reduced doubling time of ST999 in Fe^3+^ enriched conditions, the increased expression of *chuA* by ST999 in Fe^3+^ enriched conditions, and our observation that ST999 positive compartments stained highly positive for the iron probe Mem-RhoNox. However, we did not detect *chuA* in ST999 and ST95 infected BECs, possibly because haem import by ChuA is redundant at the high phagolysosomal levels of Fe^3+^ within BECs.

In contrast to ST999, ST95 not only carries systems for importing Fe^3+^ (*iroBCDEN, fyuA* and *fepA-E/G*) and haem (*chuA* and *hma*), but also *sitABCD* genes for importing Fe^2+^. This explains our finding that ST95 grows better in iron depleted conditions similar to that of the urinary tract. Indeed, the expression of *clbN* and *senB* also increased in iron depleted conditions. Also, high growth rates of UPEC in low-iron urine of UTI patients has been reported [18] along with evidence of colibactin production by UPEC strains [27]. The other two UPEC strains ST131 and ST1981 also carry both Fe^2+^ and Fe^3+^ importers, and thus might be expected to also grow better at low iron concentrations. Nevertheless, of note is that none of them carry toxins found in ST95.

Our findings that the UPEC strains survive intracellularly in BECs and, except ST95, macrophages are in line with the reported intracellular bacterial communities of UPEC. It has been reported that intracellular UPEC reservoirs are LAMP1 and LC3 positive [6]. In BECs, we observed more localization of ST999 and ST95 to LAMP1 positive vacuoles compared to XL1 blue. Moreover, both ST999 and ST95 increased LC3 levels, which might be indicative of increased autophagy flux [60]. LC3 can be associated with vacuoles due to activated xenophagy or LC3-mediated phagocytosis (LAP) [61]. As intracellular pathogenic bacteria escape from phagolysosomes, they are tagged by cytosolic ubiquitin-conjugating enzymes leading their encapsulation in double membrane vacuoles in a process called xenophagy [58]. Electron microscopy showed that, in contrast to non-pathogenic *E. coli* strains, UPEC strains reside in double-membrane vacuoles within macrophages. The presence of double membranes and recruitment of LC3 to UPEC carrying vacuoles indicate xenophagy. LC3 was also recruited to vacuoles carrying non-pathogenic XL1 blue and MG1655 strains, but, in contrast to UPEC, only a single membrane was present at these vacuoles. These findings suggest that non-pathogenic *E. coli* where cleared by LAP, as has been reported previously for bacteria [61].

We found that whereas ST95 persisted in BECs, it caused macrophage death and this might well contribute to its immune evasion. This selective macrophage death does not seem to be solely attributable to hemolytic activity, because we found that ST95 exerts this activity in both macrophages and BECs. Therefore, the different conditions within the vacuoles of macrophages and BECs likely differentially induced expression of other toxins that triggered macrophage death. One good candidate toxin for this is colibactin, a genotoxic protein that was recently found in ∼25% of UPEC strains [27] and its expression is upregulated at low iron concentrations [26]. Indeed, our data also show an increased and decreased transcription in ST95 of *clbN* (the gene coding for colibactin) in Fe^3+^ depleted and enriched conditions, respectively. Thus, colibactin expression might be repressed in the presence of the higher labile iron pool in BECs, whereas it will be expressed at the lower iron concentrations in macrophages. However, we did not manage to detect *clbN* transcripts in ST95 infected BECs and macrophages, possibly due to the limited detection sensitivity and/or because of the transient nature of *clbN* expression. Similarly, Shigella enterotoxin *senB* also showed increased expression at Fe^3+^ depleted conditions. However, likely because this iron dependency was only low (<2-fold), *senB* expression did not significantly alter between ST95 infected macrophages and BECs.

Thus, we have shown that pathogenic UPEC strains adopt different but overlapping strategies to persist in the urinary tract and escape immune clearance. We found that whereas all UPEC strains can survive within the iron rich endolysosomal compartments of macrophages, they have different requirements for iron and some also persisted in BEC. While most pathogenic ST95 grow at the low iron concentrations similar to that of the urinary tract and selectively kills macrophages, but not BECs. This cell type specific toxicity is likely induced by the synergistic action of hemolysin, colibactin and Shigella enterotoxin. In the future, we envision that genetic screening for virulence genes might enable the treatment of UTIs with strain-specific therapies, such as combinations of iron scavengers and membrane (im)permeable antibiotics.

## Supporting information

Delivery of iron-rich endosomes/lysosomes to XL1 containing vacuoles. Green: deoxygenated iron probe. Magenta: labelled XL1 blue.

Delivery of iron-rich endosomes/lysosomes to XL1 containing vacuoles. Green: deoxygenated iron probe. Magenta: labelled XL1 blue.

Cell 1 with overlay of BF channel is shown and unlabeled video.

Delivery of iron-rich endosomes/lysosomes to ST999 containing vacuoles. Magenta: labelled bacteria. Green: deoxygenated iron probe.

Delivery of iron-rich endosomes/lysosomes to ST999 containing vacuoles. Magenta: labelled bacteria. Green: deoxygenated iron probe.

Delivery of iron-rich endosomes/lysosomes to ST999 containing vacuoles. Magenta: labelled bacteria. Green: deoxygenated iron probe.

Delivery of iron-rich endosomes/lysosomes to ST999 containing vacuoles. Magenta: labelled bacteria. Green: deoxygenated iron probe.

Cell 1 with overlay of BF channel is shown and unlabeled ST999.

Fast movement of iron-rich endosomes/lysosomes, and macrophage swelling and blebbing indicative of its death by ST95. Magenta: ST95. Green: iron probe

Fast movement of iron-rich endosomes/lysosomes, and macrophage swelling and blebbing indicative of its death by ST95. Magenta: ST95. Green: iron probe

Fast movement of iron-rich endosomes/lysosomes, and macrophage swelling and blebbing indicative of its death by ST95. Magenta: ST95. Green: iron probe

Fast movement of iron-rich endosomes/lysosomes, and macrophage swelling and blebbing indicative of its death by ST95. Magenta: ST95. Green: iron probe

Cell 1 with overlay of BF channel is shown and unlabeled ST95.

Multiple ST999 within LAMP1 positive vacuoles.

## 5. Acknowledgement

GvdB has received funding from the European Research Council (ERC) under the European Union’s Horizon 2020 research and innovation program (grant agreement No. 862137). DD thank Sjors Maassen (Molecular Immunology and Microbiology, University of Groningen) for technical support for flow cytometry, Anne de Jonge (Molecular Genetics, University of Groningen) for technical advice on whole genome isolation and Beijing Genomics Institute (BGI), Denmark for whole genome sequencing.

## 6. Author contributions

DD and GvdB designed the experiments. DD wrote the paper with critical feedback from GvdB. DD isolated whole genome of UPEC strains, and HG assembled and annotated genomic data, and carried out all bioinformatics analysis. DD carried out infection assays, and embedding for electron microscopy. RdB sectioned, and carried out electron microscopy image acquisition and analysis. DD carried out all other experiments and their analysis. MdV provided UPEC strains. Masato Niwa, and Tasuku Hirayama synthesized the iron probe.

## Data availability

The reads for the assembly of the UPEC strains genomes will be deposited in the Sequence Read Archive (SRA) at the NCBI, and the whole-genome shotgun sequences will be deposited in the GenBank database.

## Figures and Tables

**Supplementary Figure 1.**
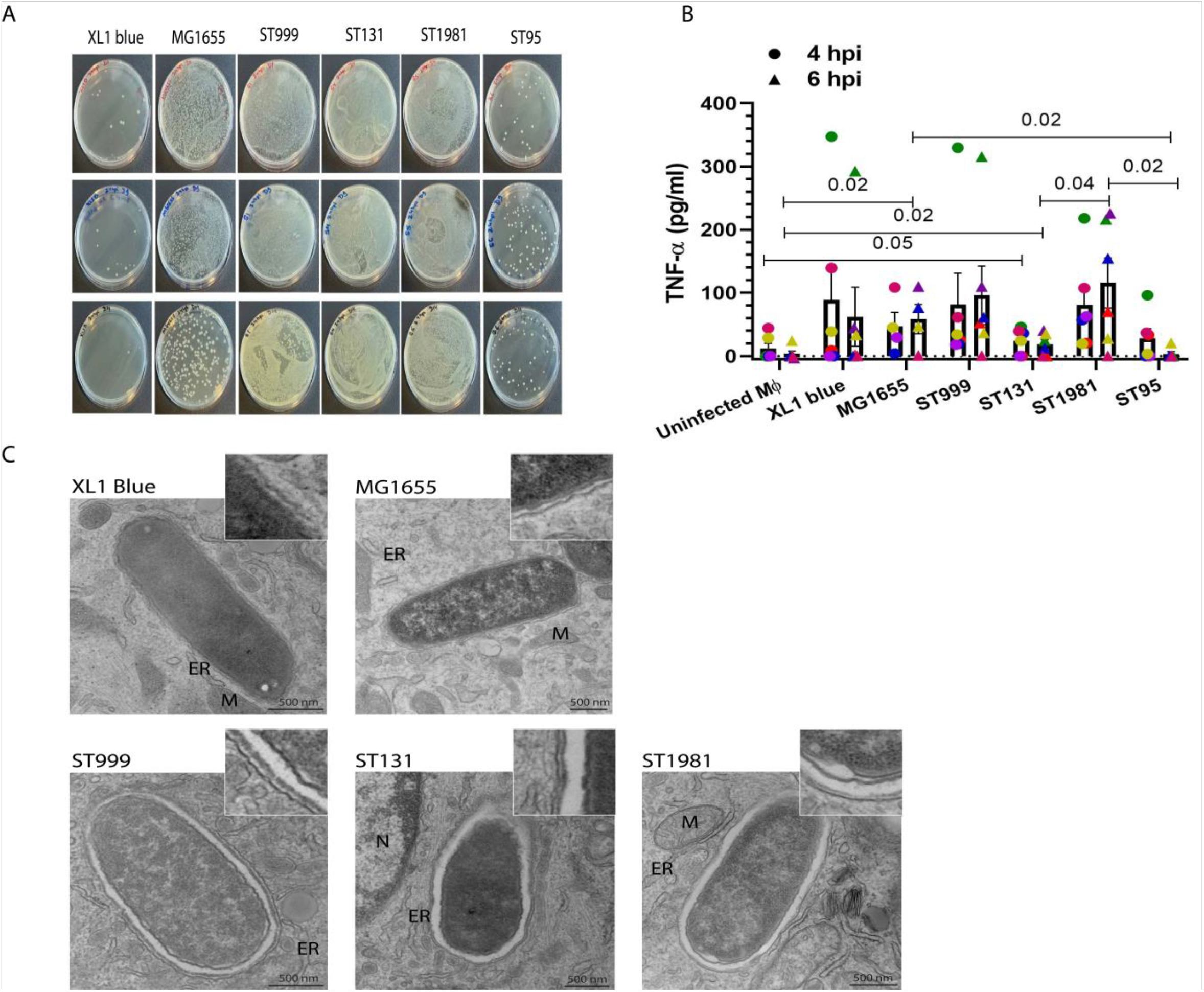
ST999, ST131, ST1981 and ST95 survive inside double membrane vacuoles in macrophages. (A) Agar plates of colony forming unit (CFU) assay showing intracellular survival of ST999, ST131, ST1981 and ST95 and XL1 blue and MG1655 at 24 hr post-infection (hpi). (n = 3 experiments). (B) TNF-α production by ELISA (n = 6 experiments, Two-tailed unpaired t-test.). (C) Representative negative staining transmission electron microscopy images showing single membrane vacuoles containing XL1 blue and MG1655 and double membrane vacuoles containing ST999, ST131 and ST1981. Scale bar, 500 nm.

**Supplementary Figure 2.**
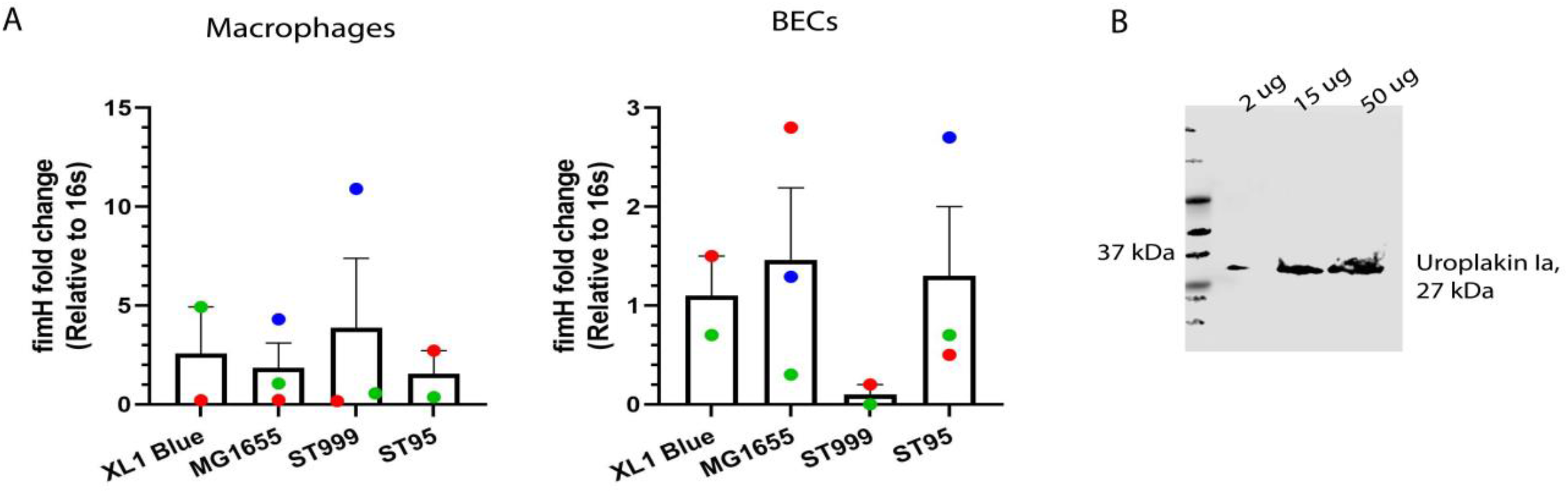
Expression of type 1 fimbriae subunit *fimH* in UPEC, and uroplakin Ia in BECs. (A) Expression of *fimH* in XL1 blue, MG1655, ST999 and ST95 isolated from infected macrophages (left) and BECs (right) at 1 hr post infection by RT-qPCR. (B) BECs were differentiated as indicated by the expression of uroplakin Ia that interacts with bacterial *FimH*. Lanes carry 2 μg, 15 μg and 50 μg protein.

**Supplementary Figure 3.**
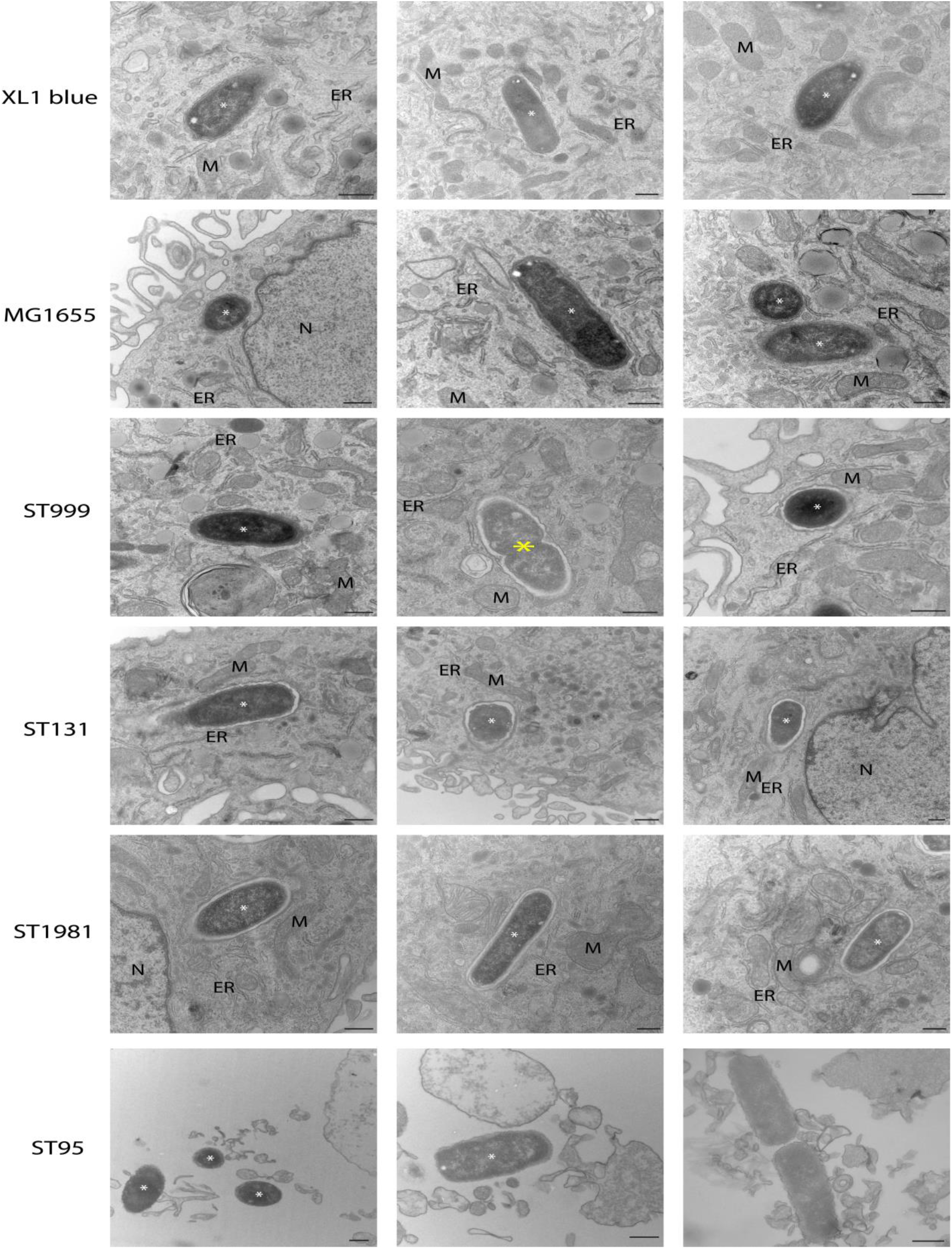
Panel of representative negatively stained transmission electron microscopy images of 1 h post infected macrophages. White asterisks: bacteria. Note: septum of ST999 suggests multiplying bacteria (yellow asterisk), and ST95 is surrounded by macrophage debris. Scale bar, 500 nm.

**Supplementary Figure 4.**
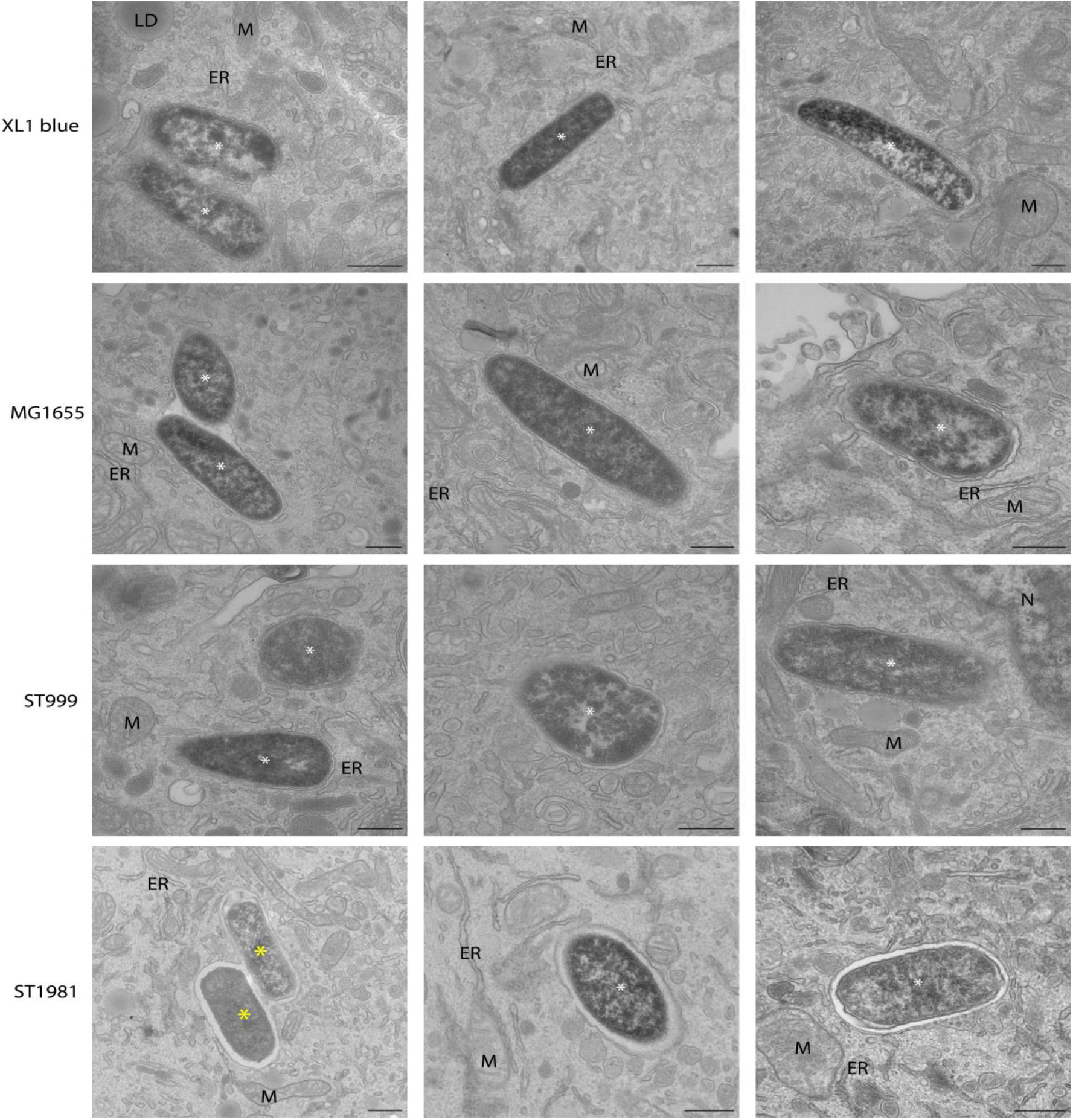
Panel of representative negatively stained transmission electron microscopy images of 4 h post infected macrophages. Note: Fused vacuoles containing ST1981 (yellow asterisk). Scale bar, 500 nm.

**Supplementary Table 1.** UPEC catogories based on ST, fimH and O-H typing. Antibiotic resistance genes, genome sizes, virulence factors and prophages is also shown.

| Samples | ST typing | Phylogroup | fimH Typing | Serotype (O-H loci) | AMR Resitance gene | Genome size | Number of VF | number of prophages (name) |
| --- | --- | --- | --- | --- | --- | --- | --- | --- |
| S1 | 999 | B2 | *fimH20* | *H4(fliC-No O gene detected)* | *mdf(A)* | 4.8 Mbp | 187 | 1 (Shigel_SfII_NC_021857(24)) |
| S4 | 131 | B2 | *fimH30* | *O25:H4(wzy-fliC)* | *aadA5*<br>*aac(3)-Ild*<br>*aac(6')-Ib-cr*<br>*mdf(A)*<br>*tet(B)*<br>*sul1*<br>*dfrA17*<br>*sitABCD*<br>*blaCTX-M-15*<br>*blaTEM-1B*<br>*blaOXA-1*<br>*qacE*<br>*catB3*<br>*catA1* | 5.3 Mbp | 194 | 4 (Entero_P88_NC_026014(22));<br>Pectob_ZF40_NC_019522(12);<br>Entero_P88_NC_026014(10);<br>Entero_BP_4795_NC_004813(7)) |
| S5 | 1981 | D | *fimH49* | *O1:H6(wzy-fliC)* | *mdf(A)*<br>*sitABCD(del)* | 5.3 Mbp | 239 | 3 (Escher_vB_EcoM_12474III_NC_049457(27);<br>Klebsi_4LV2017_NC_047818(29);<br>Entero_mEp460_NC_019716(29)) |
| S6 | 95 | B2 | *fimH18* | *O18:H7(wzy-fliC)* | *mdf(A)*,<br>*sitABCD(Peroxide)* | 5.1 Mbp | 258 | 4 (Entero_P88_NC_026014(43);Shigel_SfII_NC_021857(33);<br>Entero_mEp460_NC_019716(30);Entero_Sf101_NC_027398(13)) |
| XL1 blue | 10 | A | *fimH27* | *O16:H48(wzx-fliC)* | *mdf(A)*<br>*tet(B)* | 5.2 Mbp | 144 | 0 |
| MG1665 | 10 | A | *fimH27* | *O16:H48(wzx-fliC)* | *mdf(A)* | 4.7 Mbp | 137 | 2 (Entero_HK629_NC_019711(5);<br>Shigel_SfIV_NC_022749(8)) |

**Supplementary Table 2.**
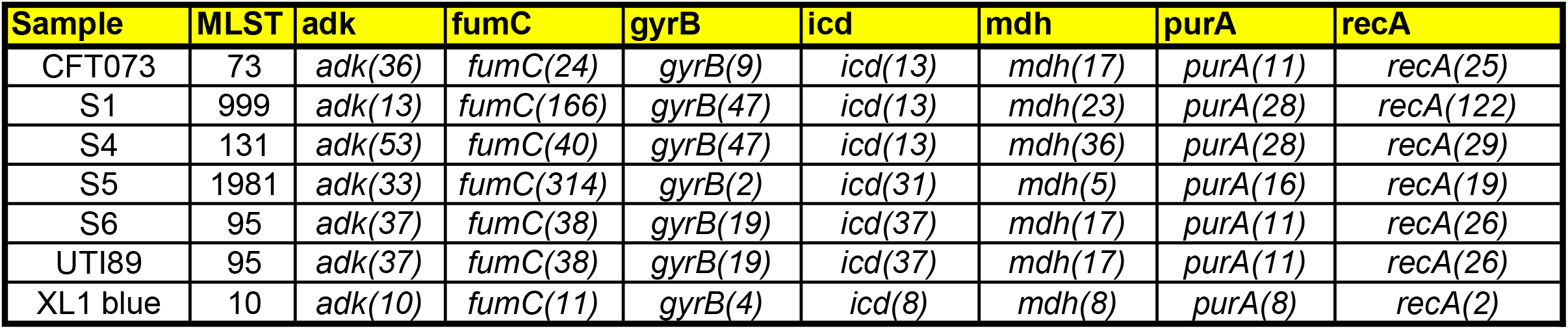
Allelic profile of housekeeping genes used for ST typing.

**Supplementary Table 3.** Comparison between ST95, UTI89 and CFT073.

| Genes | CFT073 | UTI89 | S6 (ST95) | Function |
| --- | --- | --- | --- | --- |
| *cah* | - | P | P | Divalent metal ion transporter. |
| *cnf1* | - | P | P | Toxin activates Rho GTPases RhoA, Rac1, and Cdc42 and inhibit phagocytosis. |
| *fliC* | - | P | P | component of flagella. |
| *hlyA* | P | P | P | toxin-haemolysin. |
| *hlyB* | P | P | P |  |
| *hlyC* | P | P | P |  |
| *hlyD* | P | P | P |  |
| *neuA* | - | P | P | capsular gene encodes acylneuraminate cytidyltransferase. |
| *neuC* | - | P | P |  |
| *senB* | - | - | P | Shigella enterotoxin TieB protein. |
| *sfaA* | - | P | P | S fimbriae related genes. |
| *sfaB* | - | P | P |  |
| *sfaC* | - | P | P |  |
| *sfaD* | - | P | P |  |
| *sfaE* | - | P | P |  |
| *sfaF* | - | P | P |  |
| *sfaG* | - | P | P |  |
| *sfaH* | - | P | P |  |
| *sfaS* | - | P | P |  |
| *sfaX* | - | P | P |  |
| *sfaY* | - | P | P |  |
| *clbA* | P | P | P | Colibactin - toxin, which is suggested to induce DNA |
| *clbB* | P | P | P |  |
| *clbC* | P | P | P |  |
| *clbD* | P | P | P |  |
| *clbE* | P | P | P |  |
| *clbF* | P | P | P |  |
| *clbG* | P | P | P |  |
| *clbH* | P | P | P |  |
| *clbI* | P | P | P |  |
| *clbJ* | P | P | P |  |
| *clbK* | P | P | P |  |
| *clbL* | P | P | P |  |
| *clbM* | P | P | P |  |
| *clbN* | P | P | P |  |
| *clbO* | P | P | P |  |
| *clbP* | P | P | P |  |
| *clbQ* | P | P | P |  |
| *clbR* | P | P | P |  |
| *ccbD* | - | - | P |  |
| *traT* | - | - | P | F-plasmid conjugation related |
| *traJ* | - | - | P |  |

**Supplementary Table 4.**
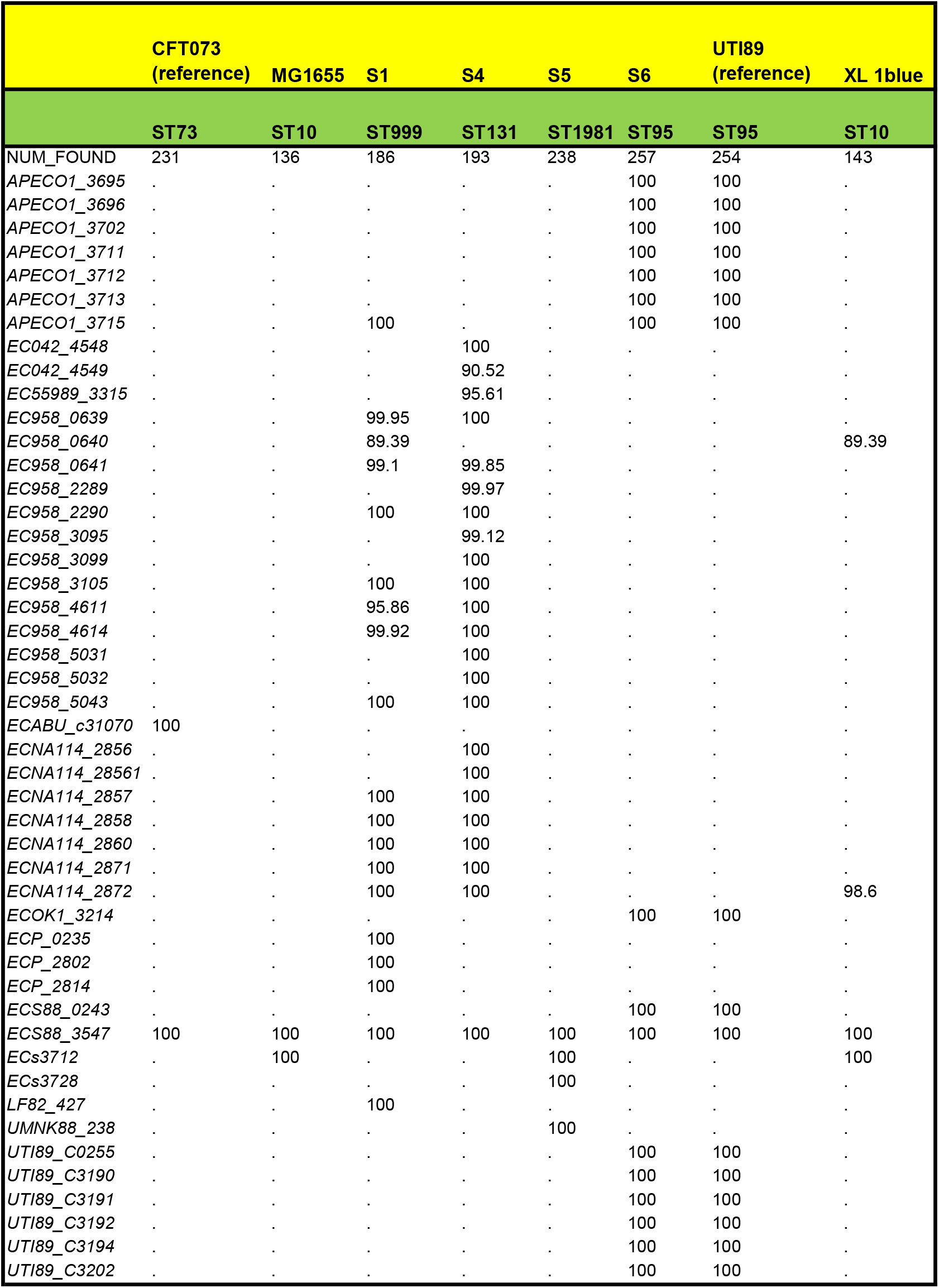
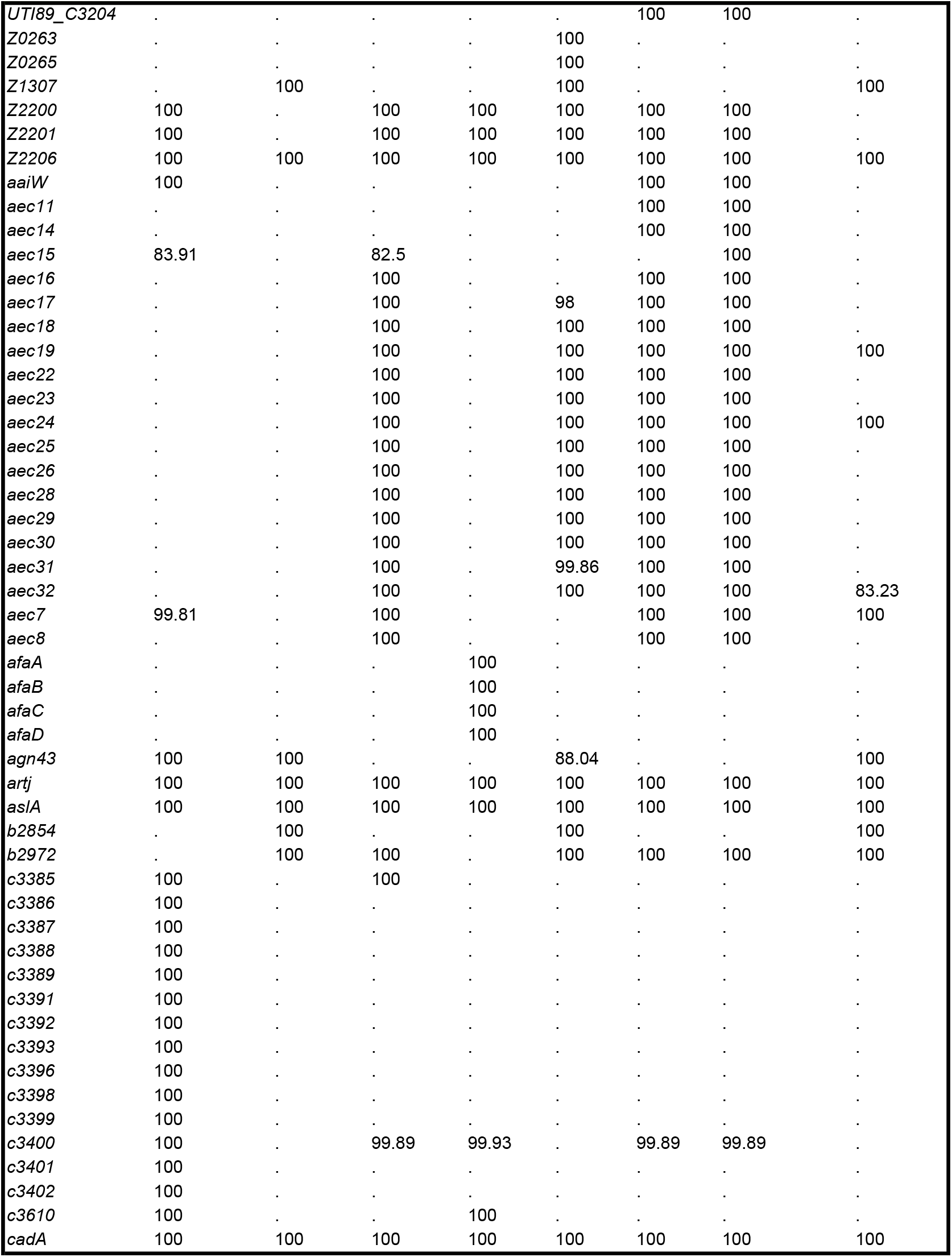
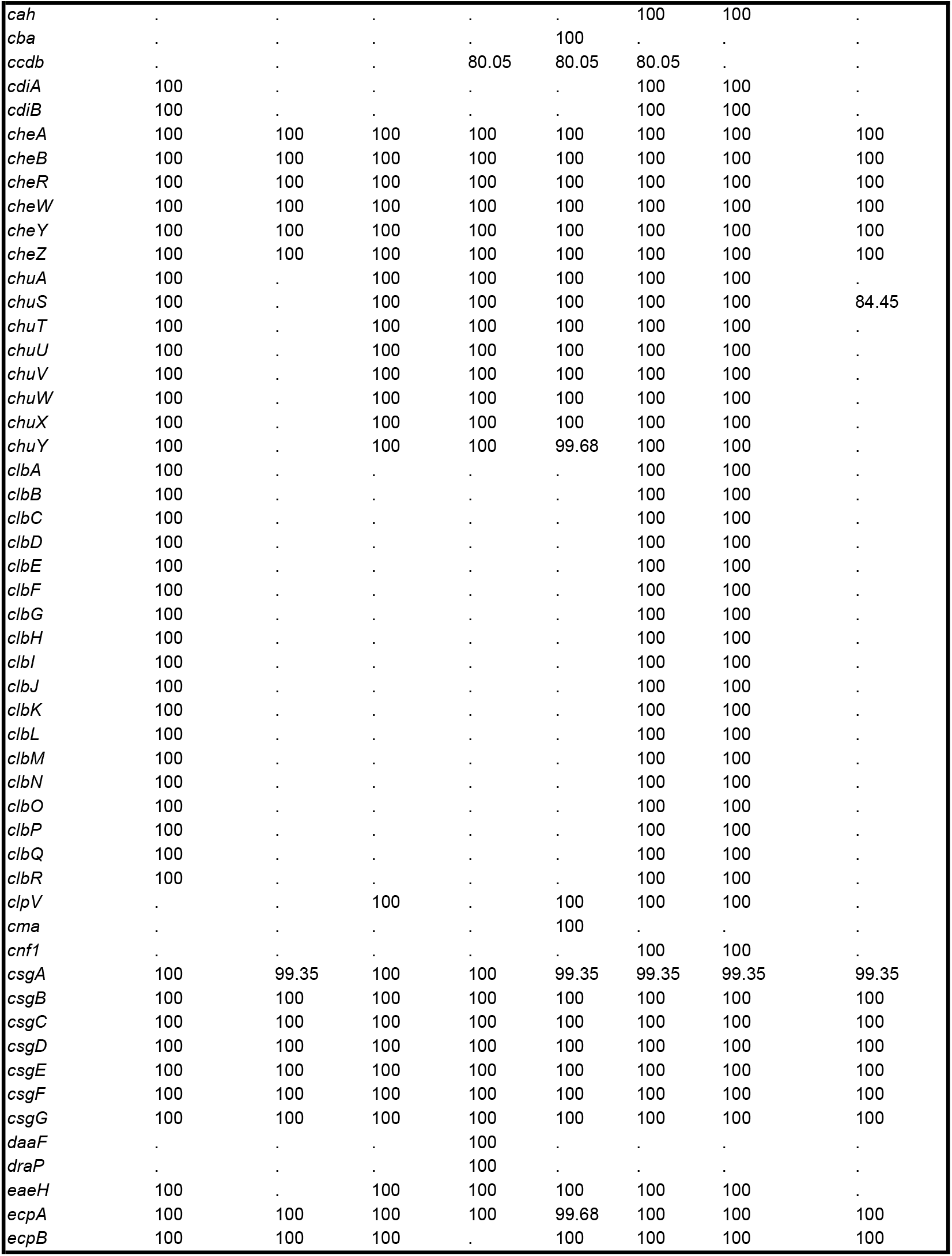
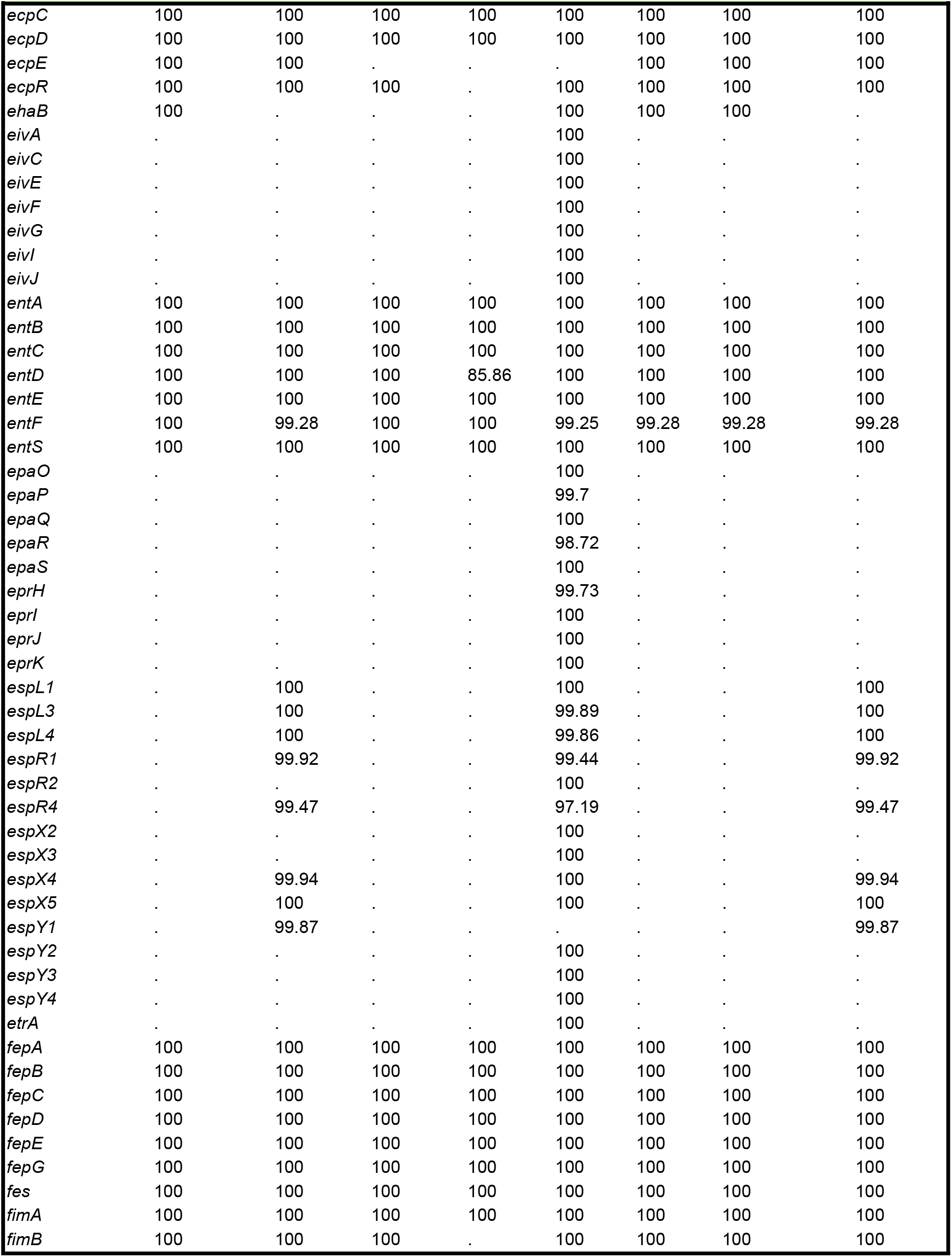
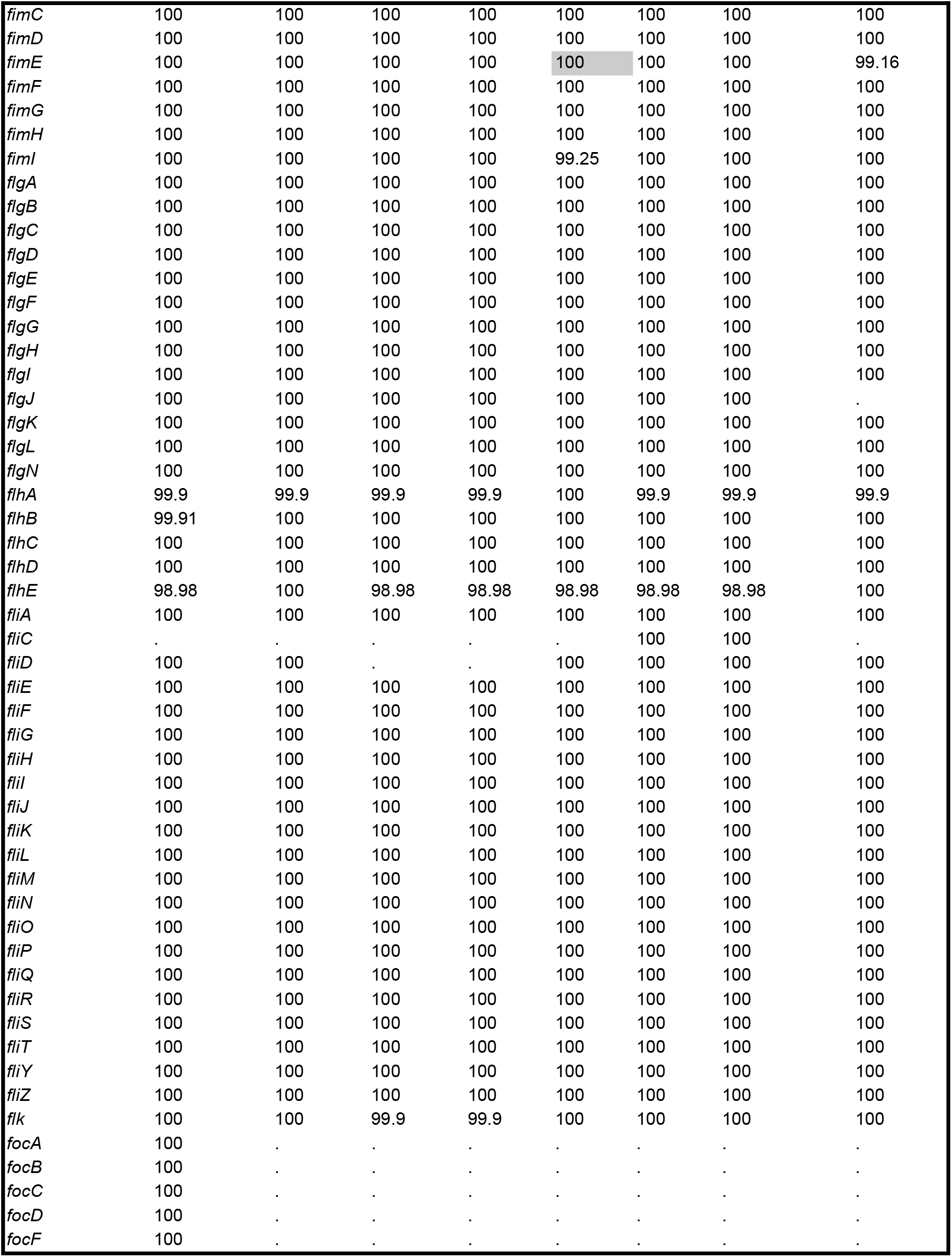
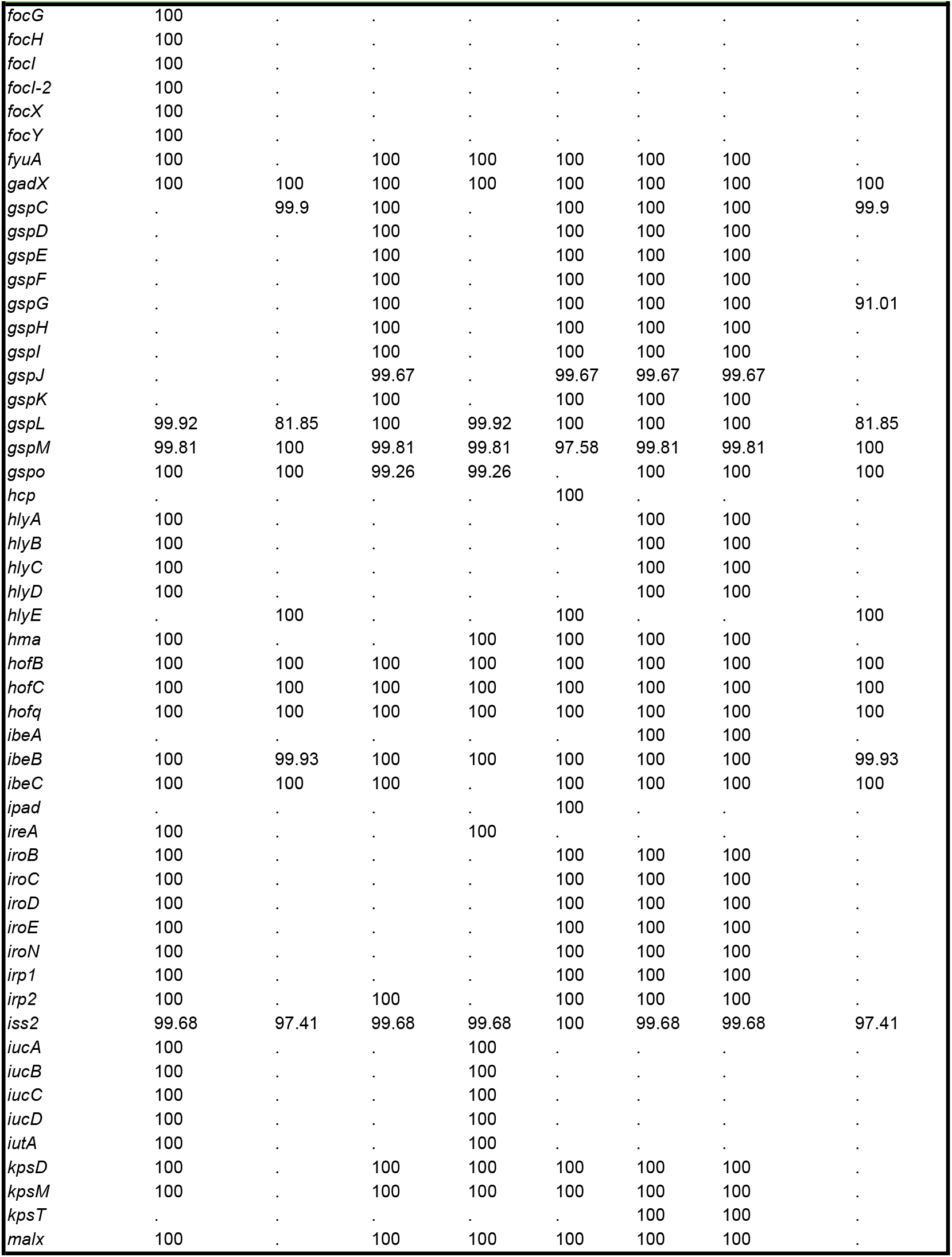
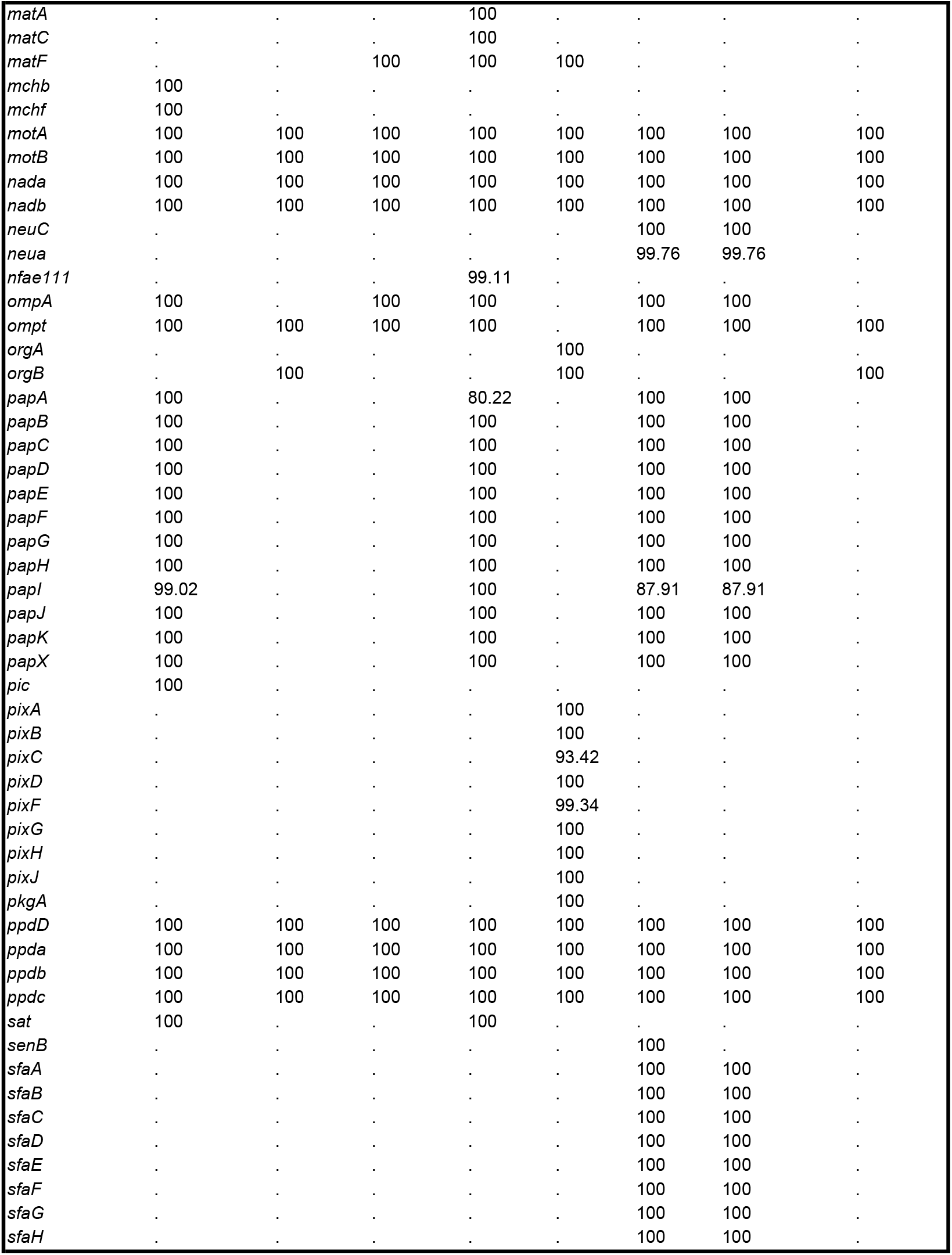
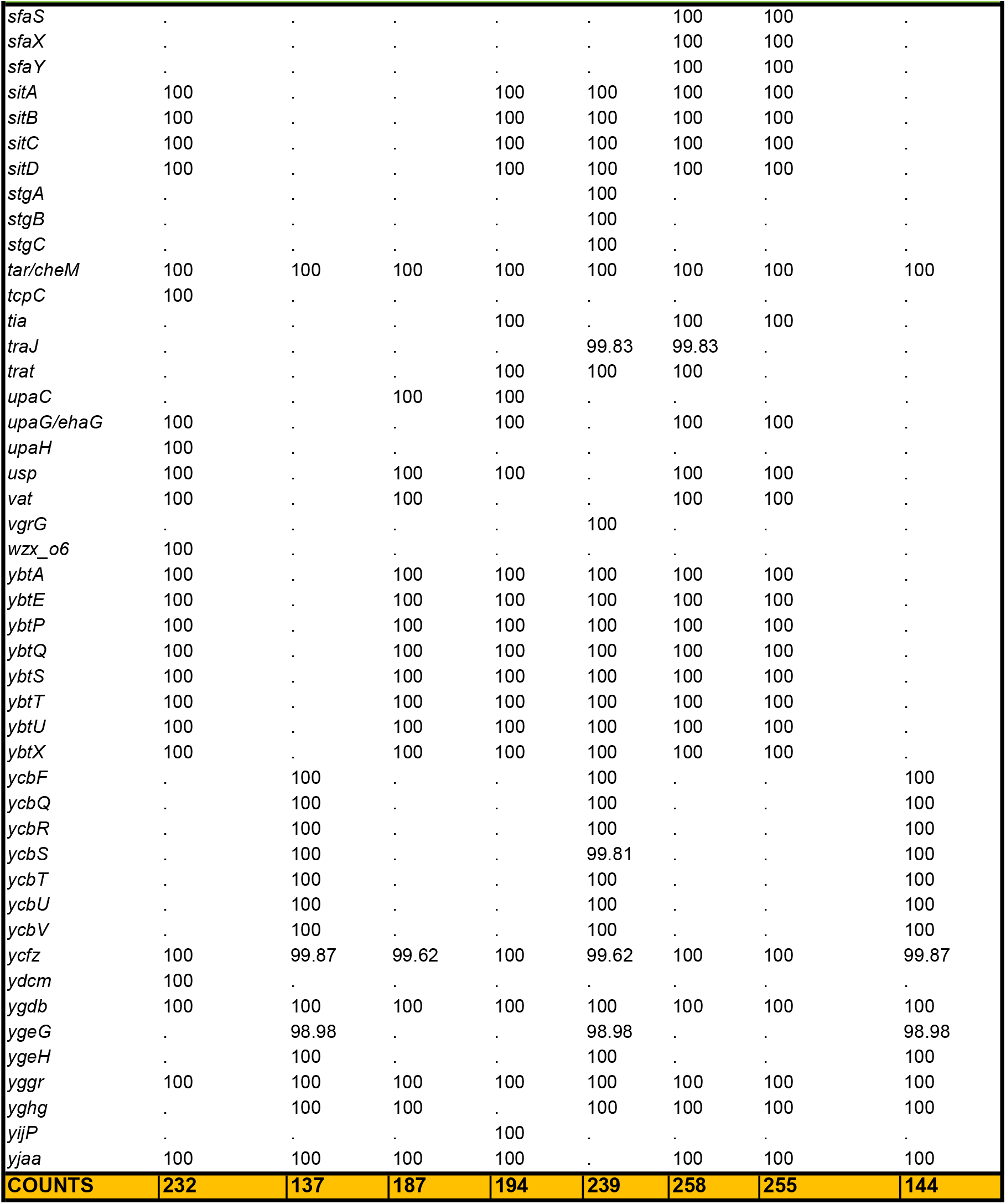
List of virulence factors.

## References

1. Moritz, R.L., R.A. Welch, The Escherichia coli argW-dsdCXA genetic island is highly variable, and E. coli K1 strains commonly possess two copies of dsdCXA. J Clin Microbiol, 2006.

2. Flores-Mireles, A.L., J.N. Walker, M. Caparon, and S.J. Hultgren, Urinary tract infections: epidemiology, mechanisms of infection and treatment options. Nature Reviews Microbiology, 2015.

3. Mann, R., D.G. Mediati, I.G. Duggin, E.J. Harry, and A.L. Bottomley, Metabolic Adaptations of Uropathogenic E. coli in the Urinary Tract. Front Cell Infect Microbiol, 2017.

4. Hayes, B.W., S.N. Abraham, Innate Immune Responses to Bladder Infection. Microbiol Spectr, 2016.

5. Olson, R.P., L.J. Harrell, and K.S. Kaye, Antibiotic resistance in urinary isolates of Escherichia coli from college women with urinary tract infections. Antimicrob Agents Chemother, 2009.

6. Dikshit, N., P. Bist, S.N. Fenlon, N.K. Pulloor, C.E.L. Chua, M.A. Scidmore, J.A. Carlyon, B.L. Tang, S.L. Chen, and B. Sukumaran, Intracellular Uropathogenic E. coli Exploits Host Rab35 for Iron Acquisition and Survival within Urinary Bladder Cells. PLOS Pathogens, 2015.

7. Dason, S., J.T. Dason, and A. Kapoor, Guidelines for the diagnosis and management of recurrent urinary tract infection in women. Can Urol Assoc J, 2011.

8. Pfrimer, K., R.F. Micheletto, J.S. Marchini, G.J. Padovan, J.C. Moriguti, and E. Ferriolli, Impact of aging on urinary excretion of iron and zinc. Nutr Metab Insights, 2014.

9. Mavromatis, C.H., N.J. Bokil, M. Totsika, A. Kakkanat, K. Schaale, C.V. Cannistraci, T. Ryu, S.A. Beatson, G.C. Ulett, M.A. Schembri, et al., The co-transcriptome of uropathogenic Escherichia coli-infected mouse macrophages reveals new insights into host-pathogen interactions. Cell Microbiol, 2015.

10. Lane, M.C., C.J. Alteri, S.N. Smith, and H.L.T. Mobley, Expression of flagella is coincident with uropathogenic Escherichia coli ascension to the upper urinary tract. Proceedings of the National Academy of Sciences, 2007.

11. Rosen, D.A., T.M. Hooton, W.E. Stamm, P.A. Humphrey, and S.J. Hultgren, Detection of Intracellular Bacterial Communities in Human Urinary Tract Infection. PLOS Medicine, 2007.

12. Mulvey, M.A., J.D. Schilling, and S.J. Hultgren, Establishment of a persistent Escherichia coli reservoir during the acute phase of a bladder infection. Infect Immun, 2001.

13. Lacerda Mariano, L., M.A. Ingersoll, Bladder resident macrophages: Mucosal sentinels. Cell Immunol, 2018.

14. Mora-Bau, G., A.M. Platt, N. van Rooijen, G.J. Randolph, M.L. Albert, and M.A. Ingersoll, Macrophages Subvert Adaptive Immunity to Urinary Tract Infection. PLoS Pathog, 2015.

15. Ingersoll, M.A., K.A. Kline, H.V. Nielsen, and S.J. Hultgren, G-CSF induction early in uropathogenic Escherichia coli infection of the urinary tract modulates host immunity. Cell Microbiol, 2008.

16. Bower, J.M., D.S. Eto, and M.A. Mulvey, Covert operations of uropathogenic Escherichia coli within the urinary tract. Traffic, 2005.

17. Haas, A., The phagosome: compartment with a license to kill. Traffic, 2007.

18. Sintsova, A., A.E. Frick-Cheng, S. Smith, A. Pirani, S. Subashchandrabose, E.S. Snitkin, and H. Mobley, Genetically diverse uropathogenic Escherichia coli adopt a common transcriptional program in patients with UTIs. Elife, 2019.

19. Wirth, T., D. Falush, R. Lan, F. Colles, P. Mensa, L.H. Wieler, H. Karch, P.R. Reeves, M.C.J. Maiden, H. Ochman, et al., Sex and virulence in Escherichia coli: an evolutionary perspective. Molecular microbiology, 2006.

20. Riley, L.W., Pandemic lineages of extraintestinal pathogenic Escherichia coli. Clinical Microbiology and Infection, 2014.

21. Kakkanat, A., M. Totsika, K. Schaale, B.L. Duell, A.W. Lo, M.-D. Phan, D.G. Moriel, S.A. Beatson, M.J. Sweet, G.C. Ulett, et al., The role of H4 flagella in Escherichia coli ST131 virulence. Scientific Reports, 2015.

22. Liu, B., A. Furevi, A.V. Perepelov, X. Guo, H. Cao, Q. Wang, P.R. Reeves, Y.A. Knirel, L. Wang, and G. Widmalm, Structure and genetics of Escherichia coli O antigens. FEMS Microbiology Reviews, 2019.

23. Garcia, E.C., A.R. Brumbaugh, and H.L.T. Mobley, Redundancy and specificity of Escherichia coli iron acquisition systems during urinary tract infection. Infect Immun, 2011.

24. Landraud, L., M. Gibert, M.R. Popoff, P. Boquet, and M. Gauthier, Expression of cnf1 by Escherichia coli J96 involves a large upstream DNA region including the hlyCABD operon, and is regulated by the RfaH protein. Mol Microbiol, 2003.

25. Davis, J.M., S.B. Rasmussen, and A.D. O’Brien, Cytotoxic necrotizing factor type 1 production by uropathogenic Escherichia coli modulates polymorphonuclear leukocyte function. Infect Immun, 2005.

26. Tronnet, S., C. Garcie, N. Rehm, U. Dobrindt, E. Oswald, and P. Martin, Iron homeostasis regulates the genotoxicity of Escherichia coli that produces colibactin. Infect Immun, 2016.

27. Chagneau, C.V., C. Massip, N. Bossuet-Greif, C. Fremez, J.P. Motta, A. Shima, C. Besson, P. Le Faouder, N. Cénac, M.P. Roth, et al., Uropathogenic E. coli induces DNA damage in the bladder. PLoS Pathog, 2021.

28. Nataro, J.P., J. Seriwatana, A. Fasano, D.R. Maneval, L.D. Guers, F. Noriega, F. Dubovsky, M.M. Levine, and J.G. Morris, Jr., Identification and cloning of a novel plasmid-encoded enterotoxin of enteroinvasive Escherichia coli and Shigella strains. Infect Immun, 1995.

29. Brannon, J.R., T.L. Dunigan, C.J. Beebout, T. Ross, M.A. Wiebe, W.S. Reynolds, and M. Hadjifrangiskou, Invasion of vaginal epithelial cells by uropathogenic Escherichia coli. Nature Communications, 2020.

30. Mavromatis, C., N.J. Bokil, M. Totsika, A. Kakkanat, K. Schaale, C.V. Cannistraci, T. Ryu, S.A. Beatson, G.C. Ulett, M.A. Schembri, et al., The co-transcriptome of uropathogenic Escherichia coli-infected mouse macrophages reveals new insights into host–pathogen interactions. Cellular Microbiology, 2015.

31. Bokil, N.J., M. Totsika, A.J. Carey, K.J. Stacey, V. Hancock, B.M. Saunders, T. Ravasi, G.C. Ulett, M.A. Schembri, and M.J. Sweet, Intramacrophage survival of uropathogenic Escherichia coli: differences between diverse clinical isolates and between mouse and human macrophages. Immunobiology, 2011.

32. Bankevich, A., S. Nurk, D. Antipov, A.A. Gurevich, M. Dvorkin, A.S. Kulikov, V.M. Lesin, S.I. Nikolenko, S. Pham, A.D. Prjibelski, et al., SPAdes: a new genome assembly algorithm and its applications to single-cell sequencing. J Comput Biol, 2012.

33. Seemann, T., Prokka: rapid prokaryotic genome annotation. Bioinformatics, 2014.

34. Jolley, K.A., M.C.J. Maiden, BIGSdb: Scalable analysis of bacterial genome variation at the population level. BMC Bioinformatics, 2010.

35. Beghain, J., A. Bridier-Nahmias, H. Le Nagard, E. Denamur, and O. Clermont, ClermonTyping: an easy-to-use and accurate in silico method for Escherichia genus strain phylotyping. Microb Genom, 2018.

36. Joensen, K.G., A.M. Tetzschner, A. Iguchi, F.M. Aarestrup, and F. Scheutz, Rapid and Easy In Silico Serotyping of Escherichia coli Isolates by Use of Whole-Genome Sequencing Data. J Clin Microbiol, 2015.

37. Arndt, D., J.R. Grant, A. Marcu, T. Sajed, A. Pon, Y. Liang, and D.S. Wishart, PHASTER: a better, faster version of the PHAST phage search tool. Nucleic Acids Res, 2016.

38. Page, A.J., C.A. Cummins, M. Hunt, V.K. Wong, S. Reuter, M.T.G. Holden, M. Fookes, D. Falush, J.A. Keane, and J. Parkhill, Roary: rapid large-scale prokaryote pan genome analysis. Bioinformatics, 2015.

39. Price, M.N., P.S. Dehal, and A.P. Arkin, FastTree 2 – Approximately Maximum-Likelihood Trees for Large Alignments. PLoS One, 2010.

40. Alikhan, N.-F., N.K. Petty, N.L. Ben Zakour, and S.A. Beatson, BLAST Ring Image Generator (BRIG): simple prokaryote genome comparisons. BMC Genomics, 2011.

41. Verboogen, D.R.J., N.H. Revelo, M. Ter Beest, and G. van den Bogaart, Interleukin-6 secretion is limited by self-signaling in endosomes. J Mol Cell Biol, 2019.

42. Niwa, M., T. Hirayama, I. Oomoto, D.O. Wang, and H. Nagasawa, Fe(II) Ion Release during Endocytotic Uptake of Iron Visualized by a Membrane-Anchoring Fe(II) Fluorescent Probe. ACS Chemical Biology, 2018.

43. Livak, K.J., T.D. Schmittgen, Analysis of Relative Gene Expression Data Using Real-Time Quantitative PCR and the 2−ΔΔCT Method. Methods, 2001.

44. Croxall, G., J. Hale, V. Weston, G. Manning, P. Cheetham, M. Achtman, and A. McNally, Molecular epidemiology of extraintestinal pathogenic Escherichia coli isolates from a regional cohort of elderly patients highlights the prevalence of ST131 strains with increased antimicrobial resistance in both community and hospital care settings. J Antimicrob Chemother, 2011.

45. Jensen, S.O., P.R. Reeves, Deletion of the Escherichia coli O14:K7 O antigen gene cluster. Can J Microbiol, 2004.

46. Sarkar, S., G.C. Ulett, M. Totsika, M.-D. Phan, and M.A. Schembri, Role of capsule and O antigen in the virulence of uropathogenic Escherichia coli. PLoS One, 2014.

47. Edgar, R., E. Bibi, MdfA, an Escherichia coli multidrug resistance protein with an extraordinarily broad spectrum of drug recognition. J Bacteriol, 1997.

48. Lewinson, O., J. Adler, G.J. Poelarends, P. Mazurkiewicz, A.J.M. Driessen, and E. Bibi, The Escherichia coli multidrug transporter MdfA catalyzes both electrogenic and electroneutral transport reactions. Proceedings of the National Academy of Sciences, 2003.

49. Michael, G.B., P. Butaye, A. Cloeckaert, and S. Schwarz, Genes and mutations conferring antimicrobial resistance in Salmonella: an update. Microbes Infect, 2006.

50. Ramirez, M.S., M.E. Tolmasky, Aminoglycoside modifying enzymes. Drug Resist Updat, 2010.

51. Mancini, S., M. Marchesi, F. Imkamp, K. Wagner, P.M. Keller, C. Quiblier, E. Bodendoerfer, P. Courvalin, and E.C. Böttger, Population-based inference of aminoglycoside resistance mechanisms in Escherichia coli. EBioMedicine, 2019.

52. Changkaew, K., F. Utrarachkij, K. Siripanichgon, C. Nakajima, O. Suthienkul, and Y. Suzuki, Characterization of antibiotic resistance in Escherichia coli isolated from shrimps and their environment. J Food Prot, 2014.

53. Will, W.R., L.S. Frost, Characterization of the opposing roles of H-NS and TraJ in transcriptional regulation of the F-plasmid tra operon. J Bacteriol, 2006.

54. Hayes, F., Toxins-Antitoxins: Plasmid Maintenance, Programmed Cell Death, and Cell Cycle Arrest. Science, 2003.

55. Cusumano, C.K., C.S. Hung, S.L. Chen, and S.J. Hultgren, Virulence plasmid harbored by uropathogenic Escherichia coli functions in acute stages of pathogenesis. Infect Immun, 2010.

56. Cunha, B.A., An infectious disease and pharmacokinetic perspective on oral antibiotic treatment of uncomplicated urinary tract infections due to multidrug-resistant Gram-negative uropathogens: the importance of urinary antibiotic concentrations and urinary pH. European Journal of Clinical Microbiology & Infectious Diseases, 2016.

57. Murthy, A.M.V., M.J. Sullivan, N.T.K. Nhu, A.W. Lo, M.-D. Phan, K.M. Peters, D. Boucher, K. Schroder, S.A. Beatson, G.C. Ulett, et al., Variation in hemolysin A expression between uropathogenic Escherichia coli isolates determines NLRP3-dependent vs. -independent macrophage cell death and host colonization. The FASEB Journal, 2019.

58. Sharma, V., S. Verma, E. Seranova, S. Sarkar, and D. Kumar, Selective Autophagy and Xenophagy in Infection and Disease. Frontiers in Cell and Developmental Biology, 2018.

59. Bah, A., I. Vergne, Macrophage Autophagy and Bacterial Infections. Frontiers in Immunology, 2017.

60. Klionsky, D.J., A.K. Abdel-Aziz, S. Abdelfatah, M. Abdellatif, A. Abdoli, S. Abel, H. Abeliovich, M.H. Abildgaard, Y.P. Abudu, A. Acevedo-Arozena, et al., Guidelines for the use and interpretation of assays for monitoring autophagy (4th edition)(1). Autophagy, 2021.

61. Lai, S.-c., R.J. Devenish, LC3-Associated Phagocytosis (LAP): Connections with Host Autophagy. Cells, 2012.

62. Birmingham, C.L., D.E. Higgins, and J.H. Brumell, Avoiding death by autophagy: interactions of Listeria monocytogenes with the macrophage autophagy system. Autophagy, 2008.

63. Birmingham, C.L., V. Canadien, N.A. Kaniuk, B.E. Steinberg, D.E. Higgins, and J.H. Brumell, Listeriolysin O allows Listeria monocytogenes replication in macrophage vacuoles. Nature, 2008.

64. Tran, T.T., C.D. Mathmann, M. Gatica-Andrades, R.F. Rollo, M. Oelker, J.K. Ljungberg, T.T.K. Nguyen, A. Zamoshnikova, L.K. Kummari, O.J.K. Wyer, et al., Inhibition of the master regulator of Listeria monocytogenes virulence enables bacterial clearance from spacious replication vacuoles in infected macrophages. PLOS Pathogens, 2022.

65. Schille, S., P. Crauwels, R. Bohn, K. Bagola, P. Walther, and G. van Zandbergen, LC3-associated phagocytosis in microbial pathogenesis. International Journal of Medical Microbiology, 2018.

66. Stempels, F.C., M.H. Janssens, M. ter Beest, R.J. Mesman, N.H. Revelo, M. Ioannidis, and G. van den Bogaart, Novel and conventional inhibitors of canonical autophagy differently affect LC3-associated phagocytosis. FEBS Letters, 2022.

67. Bishop, B.L., M.J. Duncan, J. Song, G. Li, D. Zaas, and S.N. Abraham, Cyclic AMP-regulated exocytosis of Escherichia coli from infected bladder epithelial cells. Nat Med, 2007.

68. Miao, C., M. Yu, G. Pei, Z. Ma, L. Zhang, J. Yang, J. Lv, Z.-S. Zhang, E.T. Keller, Z. Yao, et al., An infection-induced RhoB-Beclin 1-Hsp90 complex enhances clearance of uropathogenic Escherichia coli. Nature Communications, 2021.

69. Mojić, M., J.B. Pristov, D. Maksimović-Ivanić, D.R. Jones, M. Stanić, S. Mijatović, and I. Spasojević, Extracellular iron diminishes anticancer effects of vitamin C: An in vitro study. Scientific Reports, 2014.

70. Lv, H., P. Shang, The significance, trafficking and determination of labile iron in cytosol, mitochondria and lysosomes. Metallomics, 2018.

71. Tenopoulou, M., T. Kurz, P.-T. Doulias, D. Galaris, and Ulf T. Brunk, Does the calcein-AM method assay the total cellular ‘labile iron pool’ or only a fraction of it? Biochemical Journal, 2007.

